# Selection and drift determine phenotypic stasis despite genetic divergence

**DOI:** 10.1101/778282

**Authors:** François Mallard, Luke Noble, Thiago Guzella, Bruno Afonso, Charles F. Baer, Henrique Teotónio

## Abstract

Evolutionary theory suggests that phenotypic stasis is explained by natural selection and genetic drift, or by constraints imposed by mutation and recombination of standing genetic variation. We performed experimental evolution from standing genetic variation with the nematode *Caenorhabditis elegans*, measuring individual locomotion in outcrossing populations for 240 generations. We find that, in our constant environment, locomotion bias shows no directional divergence, due to both stabilizing and disruptive selection on specific combinations of component traits. Despite phenotypic stasis, the genetic variance-covariance structure between component traits shows clear divergence from the ancestral state and extensive differentiation among replicated populations facing the same environment. Analysis of mutation accumulation experiments and genome-sequenced recombinant inbred lines from the experimental populations indicates that the evolution of the genetic variance-covariance structure is independent of *de novo* mutation or major effect QTL; being instead explained by the joint action of selection and drift in generating subtle linkage disequilibrium differences between small effect QTL among replicate populations. These findings indicate that phenotypic evolution is repeatable because of selection, even if the genetic structuring of component traits within lineages is mostly contingent upon drift history.

## 2 Introduction

Forty years ago, Gould and Lewontin (1979) reminded evolutionary biologists that organisms cannot be meaningfully studied as atomized collections of independent traits, but rather that quantitative trait evolution must be considered in the context of the whole organism. Following the lead of earlier ideas from population genetics (Wright, 1931), Lande had already started to formalize what has come to be known as the “adaptive landscape” of quantitative trait evolution (Lande, 1976, 1979, 1980): the body of theory in which phenotypic evolution can be characterized, if not exactly in the context of an integrated whole organism, then at least as an arbitrarily large set of potentially non-independent traits. Since then, adaptive landscape theory has been greatly expanded, particularly to explain multivariate quantitative trait evolution between populations and species; reviewed in Arnold et al. (2001). Recent applications of adaptive landscape theory have shown that constraints imposed by mutation appear to have determined the course of fly wing evolution over a period of 40 million years (Houle et al., 2017), and of the early development of the nematode embryo over the past 100 million years (Farhadifar et al., 2015). These studies clearly demonstrate the utility of the adaptive landscape to explain evolutionary patterns over deep time and promise to resolve questions about the tempo and mode of phenotypic evolution as observed in contemporary species and in the fossil record, as or inferred from phylogenetic trees (Simpson, 1944).

One of the strongest predictions of the adaptive landscape is that populations will show phenotypic stasis under stabilizing selection (i.e., a lack of change in phenotypic mean values with time), and that, in the absence of environmental change, there will be phenotypic parallelism among populations, and eventually species, over the long-term (Charlesworth et al., 1982; Lande, 1986; Hansen and Martins, 1996; Estes and Arnold, 2007). The reason is that selection for a non-moving phenotypic optimum reduces phenotypic variance and will favor the evolution of a limited set of component trait combinations. However, in the short-term of tens to hundreds of generations, the standing genetic variation underlying trait variance – encapsulated in the additive genetic variance-covariance matrix describing how traits are genetically related and inherited across generations (the **G**-matrix) – is expected to diverge and lead to differentiation among populations facing similar ecological conditions, due to genetic drift and selection generating differences in linkage disequilibrium between quantitative trait loci (QTL; reviewed in Phillips and McGuigan (2006)). In the long-term of hundreds to thousands of generations, on the order of the effective population size, the pleiotropic nature of mutational effects and stochastic nature of mutational input is expected to direct which combinations of component traits are fixed among populations and species (Lande, 1980; Lynch and Hill, 1986; Barton, 1990).

Phenotypic stasis is perhaps the most common evolutionary pattern observed in extant and extinct species over timespans reaching 10^6^ years (Arnold, 2014). But the extent to which stasis of overlying phenotypes is compatible with the expected divergence of underlying genetic trait structure remains relatively unexplored, theoretically and empirically. The degree to which the adaptive landscape provides a realistic explanation of phenotypic evolution is still an open question.

From a theoretical perspective, a comprehensive analytic theory of multivariate trait evolution that includes the stochastic effects of finite population size on QTL remains out of reach [e.g., Barton and Turelli (1987); Turelli (1988); Vladar and Barton (2014)]. The simplified mathematical models employed to explain short- and long-term phenotypic evolution usually assume weak Gaussian selection on component traits, stability of the **G**-matrix, and independent effects of selection, mutation, recombination and drift [e.g., Hansen and Martins (1996); Estes and Arnold (2007); Arnold (2014)]. Whether selection is generally weak or Gaussian is an empirical problem; long-term stability of the **G**-matrix cannot be literally true. Simulations have been illuminating [e.g., Jones et al. (2004, 2007); Guillaume (2011); Draghi and Whitlock (2012); Chebib and Guillaume (2017)], but have necessarily explored a rather limited parameter space.

Empirically, with few exceptions, artificial selection experiments have employed truncation selection [e.g., Weber and Diggins (1990); Azevedo et al. (2002); Hine et al. (2014); Pelabon et al. (2010); Sztepanacz and Blows (2017)]. The advantage of these studies is that the form of selection is known and the environment is controlled; the limitation being that truncation selection is neither weak nor Gaussian, as assumed by the theory; but see, e.g., Robertson (1966); Turelli and Barton (1994). Alternatively, a number of studies have investigated evolution in natural populations [e.g., Grant and Grant (2002); Kruuk et al. (2002); Wilson et al. (2006); Hunt et al. (2007); Chenoweth et al. (2010); Roff and Fairbairn (2012); Stinchcombe et al. (2014); Ower et al. (2013); Maraqa et al. (2017)]. These have the advantage that the form of selection is not known *a priori* to violate the assumptions of the theory; the disadvantages being, first, that the effects of the environment cannot be controlled and, second, that the form of selection can usually only be inferred indirectly.

To test the adaptive landscape, we analysed the evolution of individual locomotion in an experiment with the nematode *Caenorhabditis elegans* spanning 240 generations. Our findings only broadly confirm the applicability of the adaptive landscape models, and call attention to the need to expand them to encompass the combined influence of selection and drift on multiple quantitative traits.

## 3 Results and Discussion

### 3.1 Experimental evolution of locomotion during 240 generations

We initially crossed 16 isolates to create a hybrid population with abundant standing genetic variation from which experimental evolution could ensue without waiting for *de novo* mutation (Teotónio et al., 2017). Replicate samples from the hybrid population were then domesticated for 140 non-overlapping generations at constant population size (N=10^4^) to an environment characterized by *ad libitum* food, constant temperature and relative humidity, and little to no spatial structure (Teotónio et al., 2012; Chelo and Teotónio, 2013; Carvalho et al., 2014a,b; Poullet et al., 2016). From one of the domesticated generation 140 populations, we then derived replicate populations and evolved them for another 100 generations in the same conditions (Theologidis et al., 2014; Noble et al., 2017, 2019). Evolution of trait means was followed throughout the entire experiment, the evolution of the **G**-matrix was studied during this last 100-generation stage after initial adaptation to the environment (see below). Though rare in nature (Teotónio et al., 2006), males were observed at high frequencies during the entire period of experimental evolution, and outcrossing was therefore likely to have been the predominant reproduction mode (Figure S1).

For each outbred population, or inbred founder, we measured the locomotion of approximately one thousand adult individuals using an established worm tracking method [Swierczek et al. (2011); Figure S2; see Materials and Methods 5.2.2] and computationally identified males and hermaphrodites (Materials and Methods 5.2.3). We categorized individual worm tracks into three states determined by activity and direction, and define locomotion bias as the proportion of individuals in a sample that are actively moving at the time of reproduction (the time at which embryos are extracted under our discrete-generation experimental evolution regime).

While of *a priori* unknown importance under our lab conditions, optimal variation in individual locomotion is generally crucial for finding food and mates, avoiding predators, and for dispersal to favorable habitats. We chose to follow locomotion as a phenotype for the following reasons: i) most studies testing the adaptive landscape have considered morphological phenotypes, of presumed direct relevance to fitness, but the theoretical framework accommodates any phenotype; ii) we could obtain an accurate characterization of the fitness effects of component trait variation by measuring essentially all individuals in large experimental populations at the time of reproduction; and iii) component traits of locomotion bias – the transition rates between forward, backward or still states – can be modelled on a common scale under general statistical assumptions.

We find that while the inbred hermaphrodite founders show great diversity in locomotion bias, the evolved populations rapidly attained an intermediate level of hermaphrodite activity (stationary around 40% of the time) half that of the founder average, while males are more vagile (10%). Neither hermaphrodite nor male locomotion bias showed obvious directional change or divergence from the ancestral state of the hybrid population during the 240 generations of experimental evolution (Figure 1). Among-replicate population variance in mean phenotypic values is similar or smaller than within-replicate error. Evolution of locomotion bias since hybridization of the founders therefore appears to be an example of phenotypic stasis.

**Figure 1:**
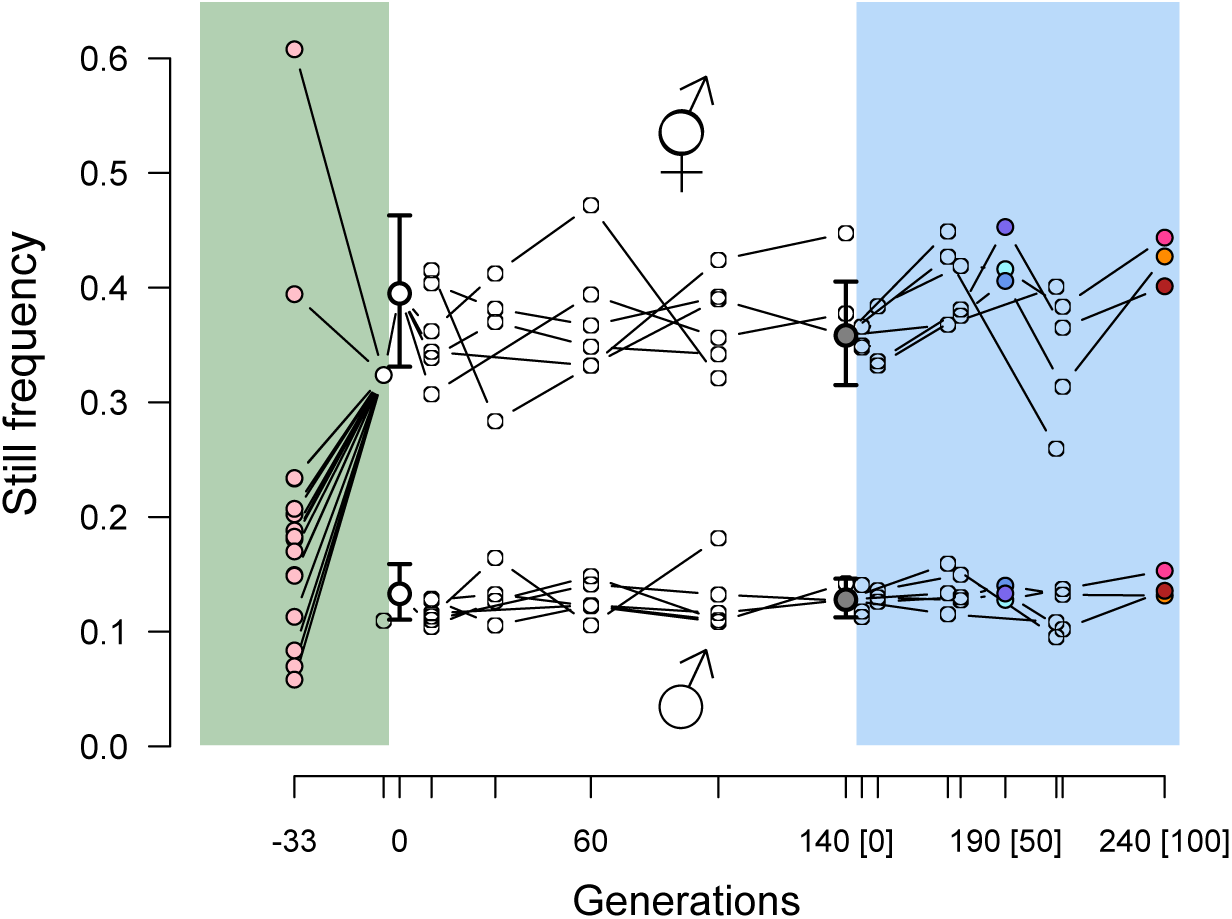
Experimental evolution of locomotion bias. The mean stationary frequency in inbred founders (wild isolates and the lab-adapted N2 reference, in pink dots) and experimental populations. Colored overlays indicate three stages of experimental evolution: hybridization (green, 33 generations of replicated pairwise crosses to generate a single ancestral population; bold-stroke white circles and 95% confidence intervals), domestication (white, 0-140 generations) and focal (blue, 141-240 generations). Axis ticks mark show sampled time points. The focal stage starts with one ancestral population (named A6140), as shown by bold-stroke grey circles and 95% confidence intervals. Point estimates are shown for 3-6 independent replicate populations at other generations. Colored points during the focal stage indicate the populations from which recombinant inbred lines were derived for **G**-matrix characterization and fitness analysis (see below) at generation 50 (named CA[1-3]50) and 100 (CA[1-3]100).

We model locomotion bias from its nine component traits, transition rates between forward movement, backward movement, and immobility, assuming spatial homogeneity and memoryless (Markov) and instantaneous temporal dynamics of individual movement (Figure 2A and Figure S3). As for locomotion bias, we see phenotypic stasis for the six independent non-self transition rates, in the sense that there is little directional change through time or divergence from ancestral states in the trait means of replicate populations, independently of sex (Figures S4 and S5). Consequently, retrospective analysis of trait means across successive replicate population samples shows that the distributions of net directional selection gradients for all transition rates are centered on zero (Figure 2B, Figure S6).

**Figure 2:**
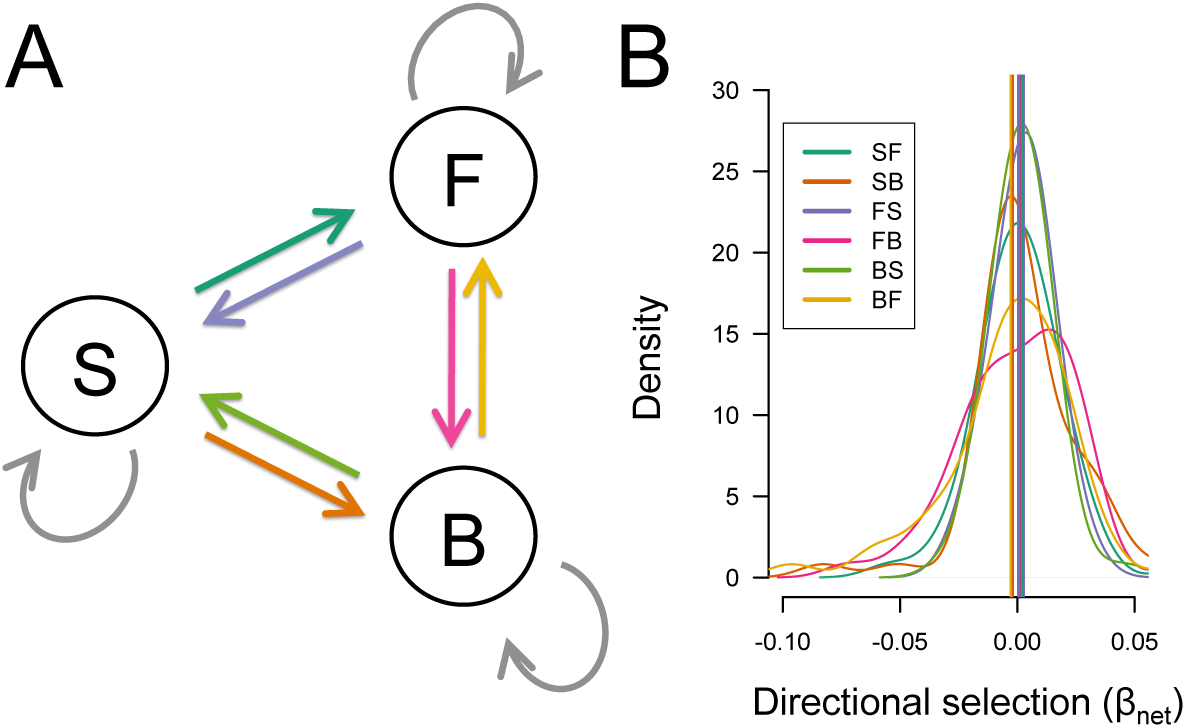
Directional selection on hermaphrodite transition rates. **A**. Component locomotion bias traits are transition rates between forward (F), backward (B) or stationary (S) states estimated from segmentation of individual worm tracking data (Methods 5.2.4). We ignore self-transition rates and consider 6 independent non-self rates shown in color. **B**. Hermaphrodite mean transition rates show little change during evolution (Figure S4). Because of this, retrospective analysis of mean transition rates assuming that the **G**-matrix is stable during the period considered indicates that there little or no directional selection (right panel, n=49 sequential time intervals, with median values indicated by vertical lines). Similar distributions of directional selection gradients on mean transition rates are found for males (Figure S6).

Given the lack of change in trait means and minimal divergence from ancestral states, locomotion appears to be neutral to our level of resolution, over this period, under our constant laboratory conditions. But stasis during 240 generations might also indicate that populations are evolving near an unknown phenotypic optimum, where stabilizing selection is common. Alternatively, locomotion might be somehow constrained by the available standing genetic variation during experimental evolution, even if the founders show that more extreme phenotypes can be realised. We next addressed these hypotheses in turn.

### 3.2 G-matrix evolution during 100 generations

We asked if the available standing variation, captured in the genetic variance-covariance structure of the six independent transition rates, could have prevented the divergence of locomotion bias after laboratory domestication. To address this, we obtained recombinant inbred lines (RILs) by selfing from the ancestral generation 140 domesticated population (approximately 200) and three replicate derived populations at generations 50 and 100 of the focal experimental stage [approximately 50 RILs for each sample; Noble et al. (2019)].

The presence and frequency of males could affect hermaphrodite locomotion (Lipton et al., 2004). Figure S7 shows little influence of male presence on hermaphrodite locomotion, since the transition rates of hermaphrodites in the inbred lines where males are not present is mostly similar to that in populations, where males were abundant. However, explicit tests for association between estimated male frequency and locomotion bias do show suggestive effects on locomotion bias and one hermaphrodite transition rate (and male frequency is positively correlated with male activity itself; see Materials and Methods 5.2.3). As discussed by Sztepanacz and Houle (2019), antagonistic selection could be constrained when genetic correlations between sexes are strong, and we cannot rule out that unmeasured evolution of sexual interactions contributes to **G**-matrix evolution. Despite this caveat, given the laboriousness of obtaining large quantities of males from each of the inbred lines (Teotónio et al., 2006), and that males and hermaphrodites showed qualitatively similar phenotypic evolution, for the remaining we focus on the evolution of hermaphrodite locomotion bias.

In the absence of net directional dominance and epistasis, the transition rate differences between inbred lines is an estimate of the population genetic variance-covariance structure in locomotion bias (**G**), the symmetric matrix where diagonal entries specify the heritable (additive) variance for each transition rate and off-diagonals the heritable (additive) pairwise covariance between transition rates. To test for inbreeding effects we compared mean trait values of the populations with those of the inbred lines derived from them (Lynch and Walsh, 1998). Inbreeding has little effect on transition rates, confirming that the **G**-matrix of the evolving populations should be reasonably well approximated by that of the inbred lines (Figure S7).

We estimated the **G**-matrices for each population during the focal stage. We find that divergence from the ancestral state and differentiation between replicate populations is extensive (Figure 3 shows a graphical pairwise comparison of trait covariances, Table S1 summarises formal comparisons of **G**-matrices using the Flury hierarchy). Eigen decomposition of the **G**-matrix yields six orthogonal combinations of transition rates, and thus the dimensions of locomotion most likely to influence evolutionary responses (Lande, 1979; Schluter, 1996). Comparing the amount and orientation of genetic variance in these dimensions reveals that the three replicate populations are significantly different from the ancestral state at generation 50 (2/3 replicates) and at generation 100 (all three replicates; Table S1). These results are independent of the statistical approach employed to test for significance of divergence and differentiation (Flury hierarchy or eigentensors; see supplementary text and figures in the Appendix).

**Figure 3:**
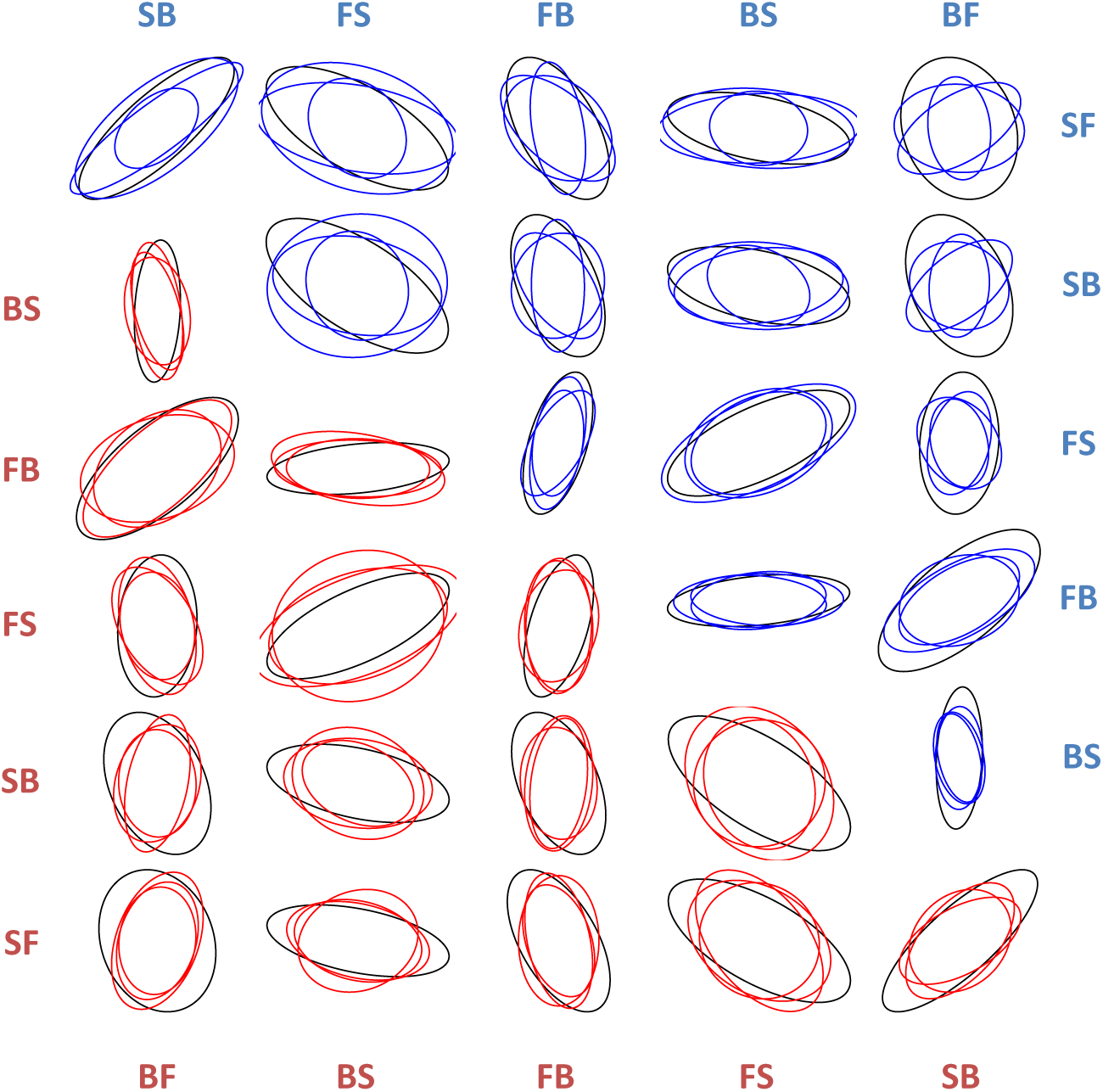
**G**-matrix divergence during 100 generations. Ellipses show the pairwise transition rate additive genetic covariances of the generation 140 domesticated ancestral population (black), and the three derived replicate populations at generation 50 (blue) and 100 (red). Ellipse width (angle) reflects the strength (sign) of covariance between traits. Ellipse axes for each trait pair (not shown) are normalised by scaling by the square-root of their eigenvalues. Significance tests of divergence from the ancestor population and differentiation among replicate populations are shown in Table S1 and as supplementary text in the Appendix.

### 3.3 G-matrix evolution by genetic drift with free recombination between QTL

**G**-matrix divergence and differentiation is expected by genetic drift alone (Lande, 1979; Lynch and Hill, 1986). Genetic drift is expected to reduce the size of **G** (variances and covariances) at a rate inversely proportional to the effective population size. Genetic drift is not, however, expected to deterministically change the average orientation of **G** among replicates, although sampling effects may contribute to the observed differentiation (Phillips et al., 2001; Whitlock et al., 2002). To test for changes in size and orientation of **G** we performed forward simulations of the generation 140 ancestral **G**-matrix during 100 generations of divergence and differentiation under drift (Materials and Methods).

Modelling a Wright-Fisher process, we considered, as during experimental evolution, that populations have an average effective population size of 10^3^ (Chelo and Teotónio, 2013) and that, for simplicity, the genomes of diploid individuals were composed of 100 freely recombining QTL, with each of the sixteen founder alleles per locus affecting up to six traits. The geno-type to phenotype map was thus modelled with variable (random) pleiotropy, although this parameter does not appear to much influence the results we obtained (analysis not shown). Starting values for QTL allele frequencies were sampled from the observed single nucleotide polymorphism (SNP) data of Noble et al. (2019) obtained for the inbred lines (Materials and Methods), and values for the **G**-matrix were based on the generation 140 ancestral population.

We find that genetic drift alone is unlikely to explain the observed divergence and differentiation of locomotion **G**-matrices. First, the main phenotypic dimension summarizes about 60% of the genetic variance in the ancestral population on average, a value that is reduced with experimental evolution beyond that seen on average under genetic drift (Figure 4A). Second, and more clearly, the orientation of the three phenotypic dimensions accounting for close to 90% of genetic variation change more, and as soon as generation 50, than that expected from 100 generations of genetic drift (Figure 4B). These results suggest that factors in tandem with genetic drift, such as selection and mutation resulted in heterogeneity in linkage disequilibrium (LD) between causal QTL among replicate populations.

**Figure 4:**
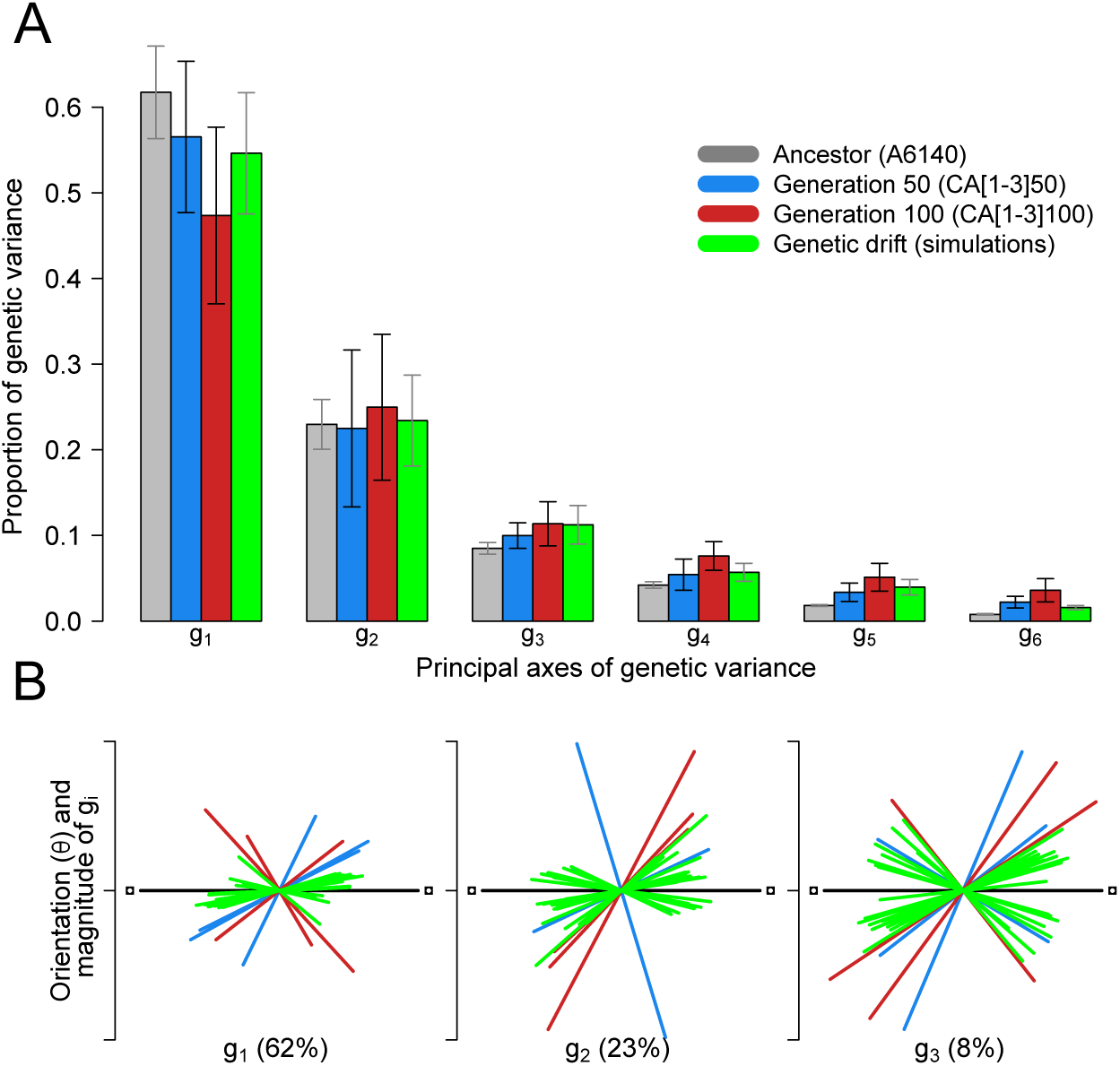
**G**-matrix evolution by genetic drift. Eigen decomposition of the 6×6 transition rate **G**-matrix yields a rotation that maximizes genetic variance in six orthogonal dimensions (principal axes, *g_i_*). The variance in these principal axes is shown in panel **A** for experimental populations (gray, blue and red) and those resulting from simulations of genetic drift (green; Materials and Methods). Grey error bars indicate the standard deviation obtained from sampling (1) 100 matrices from the posterior distribution of the experimental **G**-matrices for the ancestor population or (2) 50 simulated **G**-matrices. Black error bars are the standard deviation among the three replicate experimental populations at generation 50 and 100. **B**. The length and orientation of the first three principal axes of genetic variance among derived and simulated **G** matrices (*g_i_*), relative to those of the ancestral population arbitrarily fixed on the horizontal. The amount of genetic variance each of these axes encompasses in the ancestral population is shown in brackets.

### 3.4 Linkage disequilibrium and genetic (co)variances

Our simulations of genetic drift assumed that causal QTL were unlinked during evolution. However, genetic drift together with small differences between replicate populations in recombination rates and/or reproduction mode could lead to **G**-matrix differentiation (Lande, 1977, 1980). Effective recombination of QTL is a function of on-going meiotic crossing-over/gene conversion (Noble et al., 2017) and outcrossing rates (Figure S1), with concomitant reduction in linkage disequilibrium, which was extremely strong at the start of the hybridization stage and has been progressively eroded, on average, during domestication and the 100 generations of the focal stage (Noble et al., 2019).

Given power limited to mapping only large effect QTL for these polygenic traits (L. Noble, personal observation; manuscript in preparation), we cannot explicitly follow linkage disequilibrium between all causal variation. As an alternative, more general but indirect, approach, we examined the effect of independently varying the within-population weighting of LD on **G**-matrices estimated from genome-wide RIL SNP data (Noble et al., 2019) (see Materials and Methods 5.3.6). With this analysis we observed variability in both directional effects of LD weight on divergence from the ancestral population for some trait combinations, and also variation in the magnitude of the (co)variance differentiation at generation 50 and generation 100 (Figure 5). This suggests changes in effective recombination between causal alleles underlie at least some of the observed differentiation in **G**, and poses the question of what processes underlie this change.

**Figure 5:**
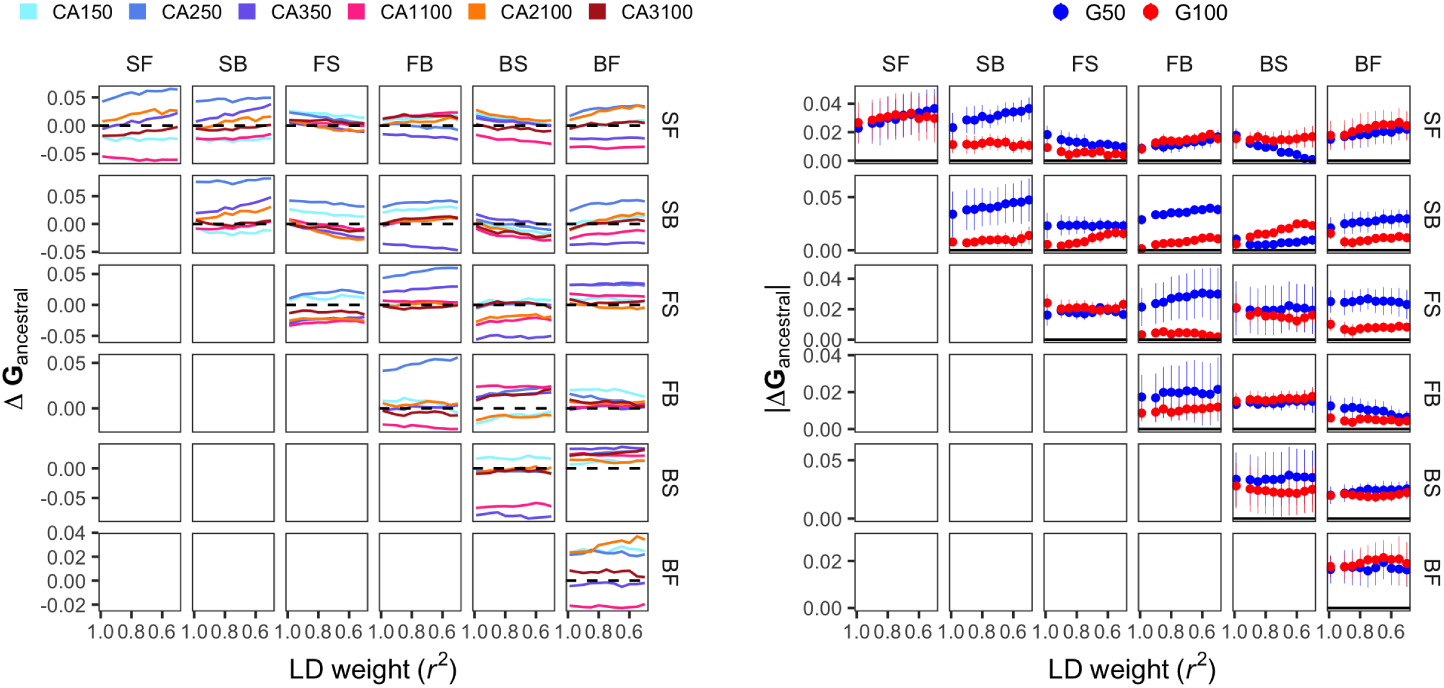
Direction and magnitude of LD effects on **G** divergence and differentiation. Here, **G** is estimated from diallelic markers genome-wide in recombinant inbred lines (RILs). LD weighting is the maximum *r*^2^ threshold for independently pruning segregating genotypes in each population before construction of an additive genomic relationship matrix (**A**) and estimation of **G**. Left: deviations in trait (co)variances from the ancestor as a function of LD weight, where each point shows the value for a given population relative to **A** based on all markers, with a value of 0 indicating equivalence with the ancestral population. Consistent directional effects of LD on divergence are apparent for some trait covariances (e.g., BS/SB), while variation in the strength (slope) and direction of covariance could contribute to differentiation of **G**. Right: the mean absolute deviation (with SE) in LD effect also varies between G50 and G100 (e.g., FB/SB, BS/SB).

### 3.5 The role of mutation

It is possible that in the short-term of our experimental scale, mutational input could contribute to heterogeneity of **G**-matrices among replicate populations due to limited stochastic sampling of mutation. To address the role of mutation we measured locomotion bias in more than 120 inbred lines that accumulated mutations in a near-neutral fashion for 250 generations (Baer et al., 2005; Yeh et al., 2017) (Methods 5.3.3). These mutation accumulation (MA) lines were derived from two of the 16 founders of our hybrid population (N2 and PB306).

As for the standing genetic variation described by **G**-matrices, the structure of the mutational genetic variance-covariances between inbred lines (**M**-matrices) is informative as to how much of the divergence and differentiation in transition rates during experimental evolution can be generated by mutation. We found that while new mutations could have generated a sufficient amount of genetic (co)variance during the 100 generations of the focal experimental stage (Figure 6; and see below), the orientations of **M**- and **G**-matrices of the three replicate populations are not aligned.

**Figure 6:**
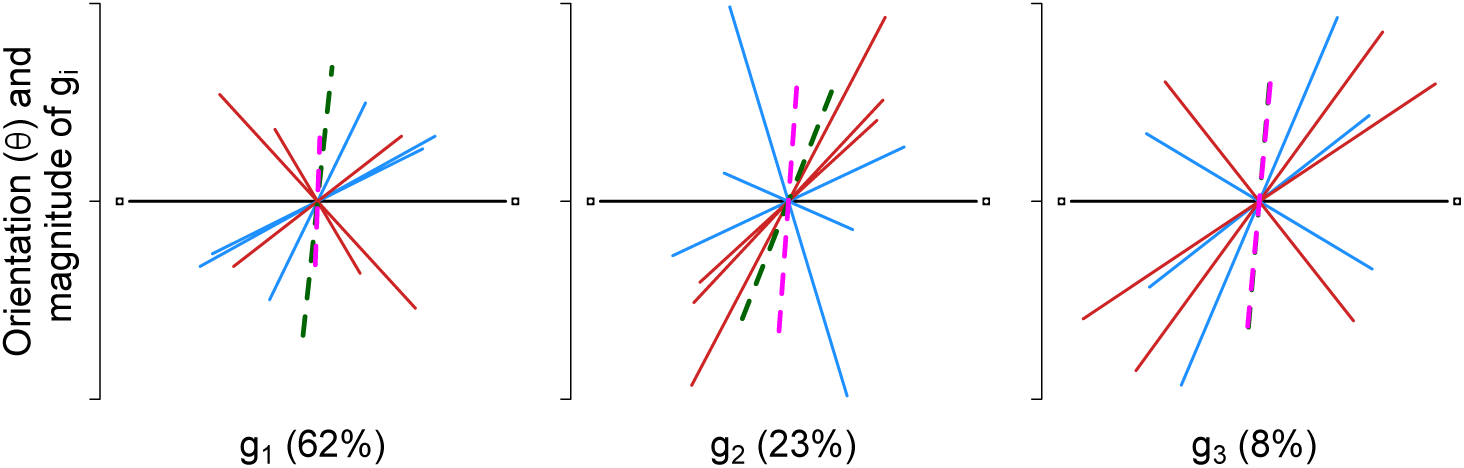
Expected mutational variance for locomotion bias after 100 generations. The length and orientation of the top three orthogonal axes, accounting for 91% and 83% of the variance across the two founder backgrounds (N2 in dark green, PB306 in pink). The **G**-matrices of experimental evolution populations are also shown for comparison (blue for generation 50, red for generation 100 replicate populations) relative, for each principal axis, to the 140 generation domesticated population **G**-matrix on the horizontal. The length of the lines indicates how much variation each axis accounts for, the orientation the particular combination of transition rates maximizing genetic variation. The proportion of variance explained by each dimension in the A6140 population is shown below each panel (from Figure 4). N2 and PB306 are aligned in *g*_3_.

Mutation could have generated sufficient genetic (co)variance for experimental evolution. However, measurement of fitness components, such as fertility and viability, in the MA lines has shown that most new mutations are, as expected, deleterious (Baer et al., 2005; Yeh et al., 2017) and are therefore unlikely to have reached appreciable frequencies during the focal 100 generations (Chelo et al., 2013b). Rare beneficial mutations are also not expected to reach high frequencies before a few hundred generations, but see Azevedo et al. (2002), with initial evolution occurring first through the sorting of standing genetic variation (Hill, 1982; Matuszewski et al., 2015). For neutral mutations, we conducted forward population genetic simulations varying the expected mutation target size (mutation rate) and modelling the population sizes realized during experimental evolution (Materials and Methods 5.3.5). These simulations show that no neutral mutation would reach sufficiently high frequency to be of much consequence during experimental evolution, even when considering very large targets/high mutation rates (Figure S8). Overall, then, mutation does not appear to have much contributed to the observed divergence and differentiation of the **G**-matrix during 100 generations of experimental evolution.

### 3.6 Quadratic selection and genetic drift

If genetic drift, alone or together with mutational bias, does not explain subtle LD differences between replicate populations underlying **G**-matrix evolution during the focal stage, then these differences can only be attributed to natural selection.

In Noble et al. (2017) we reported the fertility of many of the inbred lines employed to estimate the **G**-matrices (mostly from the ancestral domesticated population A6140). With this data we can estimate the strength and kind of selection on locomotion bias in our constant laboratory environment by multiple partial regression of the mean fertility of inbred lines on their mean transition rates (Materials and Methods 5.4.2). The partial regression coefficients obtained are: linear fitness effects of individual variation in transition rates, interpreted as revealing directional selection; quadratic fitness effects of individual variation in transition rates, if positive (negative) pointing to disruptive (stabilizing) selection on each transition rate; and quadratic effects of correlated selection between pairs of transition rates (Lande and Arnold, 1983). Together the regression coefficients describe the curvature and tilt of the multivariate “selection surface” in our environment, as summarized in part by the symmetric *γ*-matrix, where single trait coefficients are represented on the diagonal and among trait interaction coefficients on the off-diagonal. With this approach, however, we likely underestimate selection strength because fertility is not the only component of fitness (Chelo et al., 2013a, 2019). A second caveat with this approach is that we might misinterpret the effects of selection on phenotypic variance since selection differentials/gradients estimated away from the phenotypic optimum do not necessarily equate with the covariance between the individual squared deviations from the population mean with relative fitness (L.-M. Chevin, pers. communication 08/2019), as expected from evolutionary theory (Robertson, 1966; Morrissey and Bonnet, 2019).

With these caveats in mind, we find disruptive selection on backward-still (BS) and forward-still (FS) transition rates, positive correlated selection between FS and BS, positive correlated selection between SB and FS, and negative correlated selection between SF and FS transition rates (Figure S9). These results are largely insensitive to the inclusion of linear effects (two of which are significant in the full model), and we ignore them when considering the effects of quadratic selection (Figure S10, see also Figure 2).

To visualize the selection surface we decomposed the *γ*-matrix by canonical analysis (Materials and Methods 5.4.3) so that selection is interpreted only as stabilizing or disruptive on new phenotypic (canonical) dimensions of transition rate combinations (Phillips and Arnold, 1989). This analysis indicates that the 6-dimensional selection surface has an unstable equilibrium and is shaped as a saddle across the two phenotypic dimensions where selection is strongest (Figure 7). One of the principal canonical dimensions is negative, revealing stabilizing selection where individuals deviating from the optimum phenotype have lower fitness, while the other is positive, revealing disruptive selection where individuals at the extremes of the phenotype distribution have the highest fitness. In the other four phenotypic dimensions selection is weak to nonexistent.

**Figure 7:**
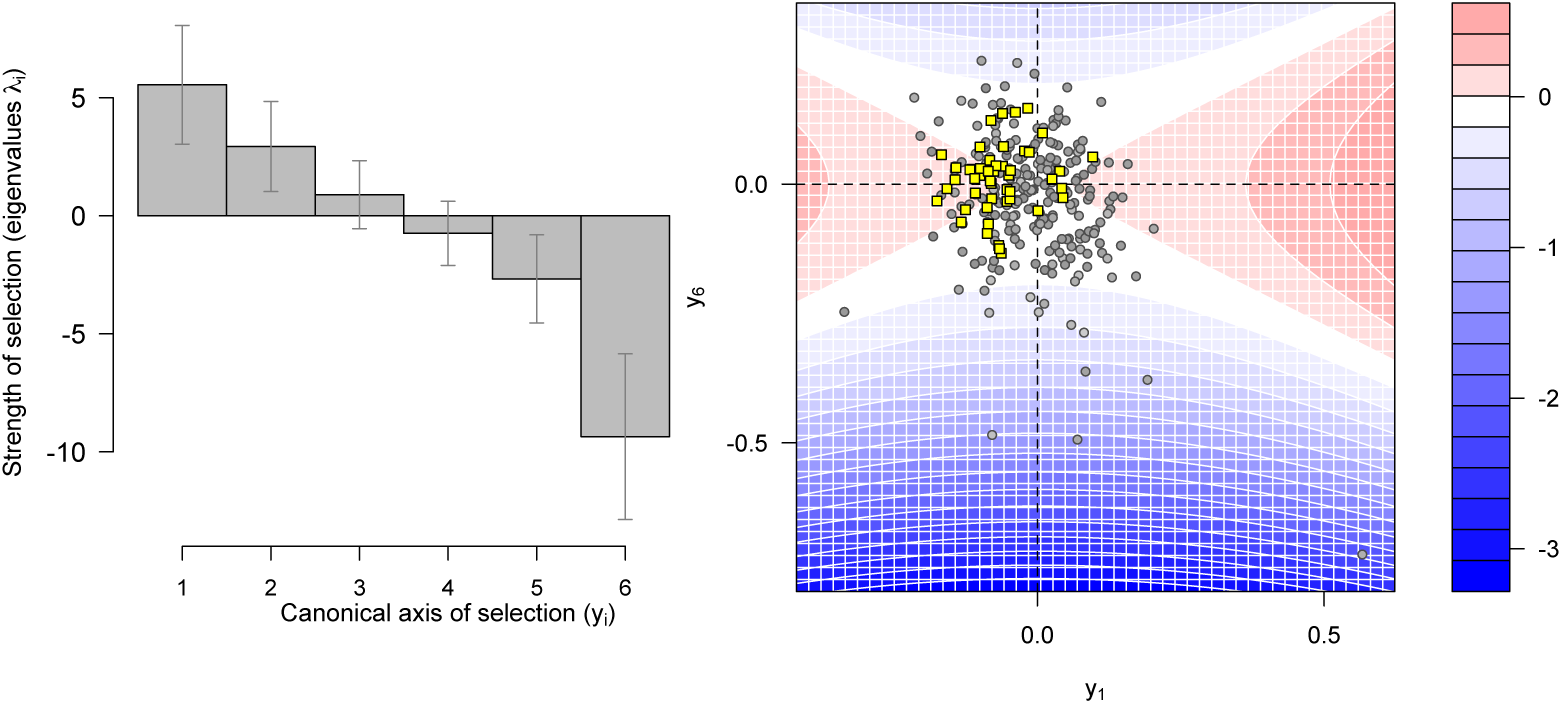
Quadratic selection and phenotypic stasis. **A**. Canonical analysis of the *γ*-matrix reveals at least one positive (*y*_1_) and one negative (*y*_6_) dimensions of transition rate combinations where selection is strong (as measured by the eigenvalue *λ*). Error bars show 95% credible intervals (1.96\**σ*, with *σ* estimated from decomposition of 5000 matrices sampled from the posterior distribution of *γ*). **B**. Selection surface for projection of transition rates onto the two main canonical axes of selection (*y*_1_ and *y*_6_ in **A**). The color code for relative fitness is centered on the stationary point (*w*_0_) and computed using equation 33 with *y*_[2_*_−_*_5]_ set to 0. Population trait means are shown as yellow squares, means of the inbred lines used to estimate the selection surface are shown as gray circles. The offset of population means from the origin is likely due to small but significant inbreeding and/or sexual interactions (Figure S7). Nevertheless, population means are within the phenotypic space probed by the inbred lines used in the inference of selection, indicating that phenotypic stasis is at least in part determined by selection. Population trait means do not evolve towards higher fitness (red) presumably because of directional selection (see main text).

Projection of mean transition rates observed during the 240 generations of domestication and focal experimental evolution onto the two main axes of the selection surface shows that phenotypic stasis may be explained by stabilizing and disruptive selection (Figure 7B). However, projection of **G**-matrices for the 100 focal generations (Materials and Methods 5.4.3) shows a clear negative relationship between the amount of additive genetic variance and selection strength that diminishes during evolution (Figure 8). This analysis therefore shows that: i) there is little opportunity for phenotypic evolution where selection is strongest, as genetic variance is lowest in these dimensions, and ii) that drift, conditional on selection, is driving genetic divergence and differentiation of the replicate populations, since a reduction in genetic variance is observed only in the phenotypic dimensions where there is weak or no selection. In the Appendix we further discuss, with eigentensor analysis of the **G**-matrices, the relative role of selection and drift in genetic divergence and differentiation of replicate populations.

**Figure 8:**
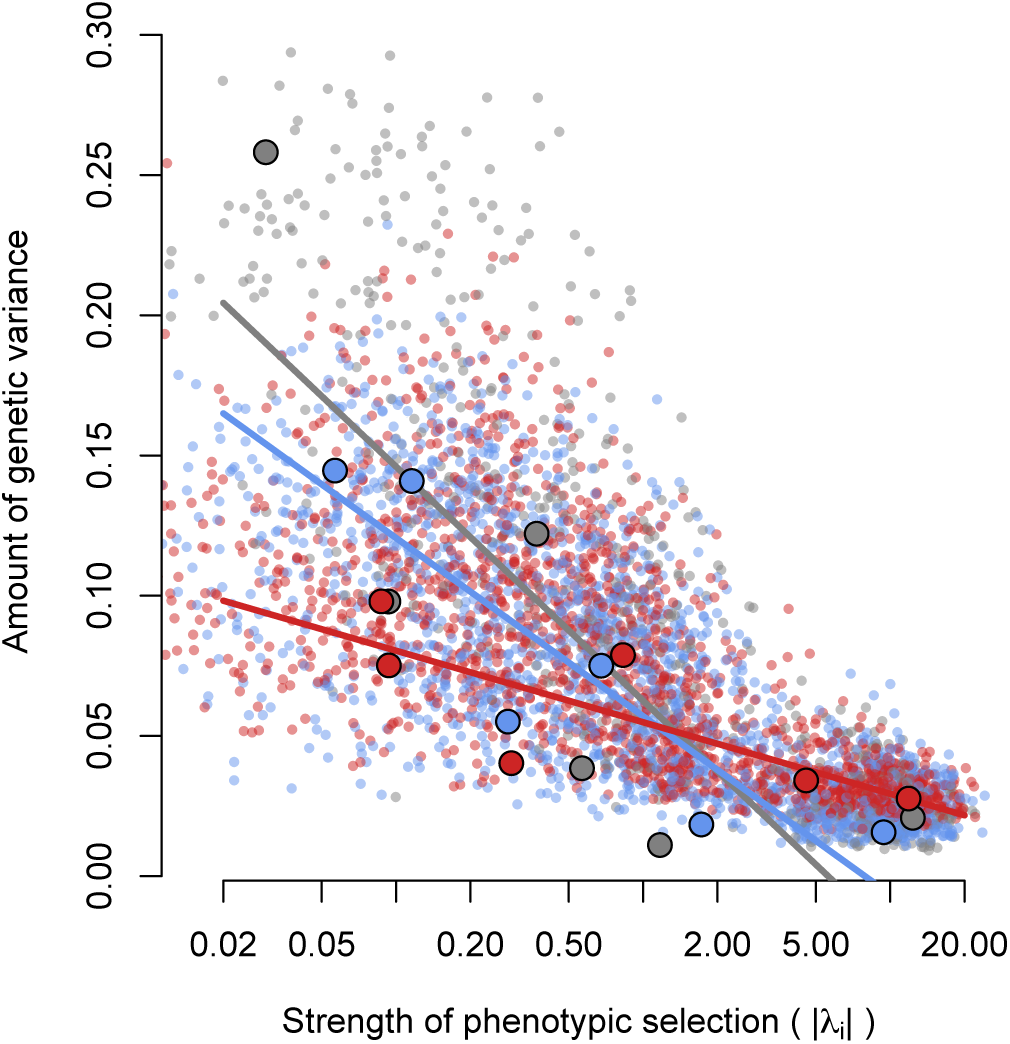
**G**-matrices align with the selection surface. Regression of the amount of genetic variance along each canonical axis on the strength of selection, estimated as the absolute value of the *γ* eigenvalues *λ_i_* (from Figure 7A). The amount of genetic variance along each canonical axis is the sum of the absolute variance-covariances along each rotated trait (*y_i_*). Small points are 100 samples from the posterior distributions of both the *γ* and **G**-matrices for each of the 7 experimental populations, with the ancestral population in grey, and evolved populations at generation 50 and 100 in blue and red respectively. Large points show the posterior mode of these samples, and corresponding lines the linear regression of genetic variance on log-transformed selection strength modes.

### 3.7 Major effect QTL

While the **G**-matrices during experimental evolution are aligned with the selection surface, this is not so for natural variation prior to experimental evolution (Figure S11). Notably, the weak negative relation between genetic variance and selection strength in the founders is due almost entirely to the presence of N2 (and the very closely related line CB4507), the *C. elegans* reference strain that has adapted to lab conditions for perhaps thousands of generations (McGrath et al., 2011; Sterken et al., 2015). While surely some N2 alleles are involved in the experimental evolution of locomotion bias (and indeed N2 haplotypes are somewhat over-represented on two of the six chromosomes), they cannot fully explain it because experimental genomes are generally well mixed (Noble et al., 2017).

To illustrate the contribution of major effect alleles from the N2 founder to the evolution of locomotion bias, we phenotyped the CX12311 near iso-genic line (NIL) that carries introgressions of the ancestral alleles of *npr-1* and *glb-5*, loci with major developmental and physiological effects located on separate chromosomes, into the N2 background (Large et al., 2016). Non-N2 *npr-1* and *glb-5* alleles are sufficient to explain a major shift in locomotion bias (Figures S12), but this occurred during the initial construction of the hybrid population and domestication, concomitant with fixation of N2 alleles in the vicinity of *npr-1* (Teotónio et al., 2012; Noble et al., 2017). The ongoing selective value of *glb-5* is unclear: after an initial large rise in frequency, there was little subsequent change during the focal stage of evolution (Figure S13). Whether due to balancing selection, or shifting effects of genetic background rendering the N2 allele increasingly neutral, *glb-5* does not appear to have been a locus of major, sustained effect under our conditions.

We also note that the lack of relationship between mutational variance and selection strength is independent of the MA background (N2, or the wild isolate PB306), a result indicating that thousands of generations of adaptation to general lab conditions are insufficient to align the **M**-matrix with the selection surface imposed by our specific environmental conditions [Figure S14; cf., Hermisson et al. (2003); Jones et al. (2007, 2014)].

After 140 generations of pre-adaptation to our conditions, the observed phenotypic stasis and differentiation of the **G**-matrix must have occurred through alleles of small to moderate additive effect (Chelo and Teotónio, 2013; Noble et al., 2017). Congruent with this expectation, all transition rate traits show significant repeatabilities – an upper bound on broad-sense heritability [Figure S15; Lynch and Walsh (1998)] – and detectable large-effect QTL explain much less than half of the variance in these traits (L. Noble, personal observation; manuscript in preparation), so the assumption of polygenicity underlying **G**-matrix theory should be reasonable (Materials and Methods 5.2.7). Overall, we conclude that both phenotypic stasis and **G**-matrix evolution are determined by selection together with drift during experimental evolution.

### 3.8 Phenotypic stasis with genetic divergence

Adaptive landscape models explaining little divergence and differentiation between populations and species rely on the assumption that there is an “average” or stable **G**-matrix and that there is stabilizing selection on a multivariate trait optimum with genetic drift. Despite the assumption of stable **G**-matrices, these models have been remarkably successful at explaining phenotypic stasis over short and long evolutionary periods (Estes and Arnold, 2007; Arnold, 2014). We finish this report by showing that these models cannot explain the phenotypic stasis we observe over the microevolutionary scale of experimental evolution, even when the assumption of a stable **G**-matrix is met.

With weak Gaussian selection (Lande, 1979), meaning that directional selection gradients are close to zero (Figure 2, Figure S10) and that the inverse of the *γ*-matrix is much smaller than the phenotypic variances observed – which is not the case during our experiment – the evolution of trait means is analogous to an Ornstein–Uhlenbeck (OU) process (Felsenstein, 1988; Hansen and Martins, 1996). We conducted simulations of this process parameterizing it with our inferred selection surface and effective population sizes, and assuming that the **G**-matrix of the domesticated 140 generation population is representative and stable (Materials and Methods 5.5). Indeed the simulations show that the evolution of canonical trait means follows a OU process (Figure S16). In the phenotypic dimension of locomotion bias under stabilizing selection there is less divergence in trait means than expected with drift, because populations randomly moving away from the optimum are pulled back by selection. In the phenotypic dimension where selection is disruptive, there is more divergence than that expected with drift, as populations randomly moving away from the fitness optimum are repelled.

However, the OU process does not explain stasis of the population mean values in locomotion bias, i.e. the frequency of active individuals (Figure 9), or in its component traits (Figure S17). First, divergence of mean trait values under OU is common, a result that is not congruent with our observations during experimental evolution (compare Figure 9 with Figure 1). Second, simulations of selection and genetic drift show similar patterns as those with only drift, a result that is not consistent with the observed quadratic selection coefficients (compare Figure 9 with Figure 8). Results are qualitatively similar irrespective of the **G**-matrix that is assumed to be stable, although the precise levels of divergence do depend on the chosen **G** (not shown).

**Figure 9:**
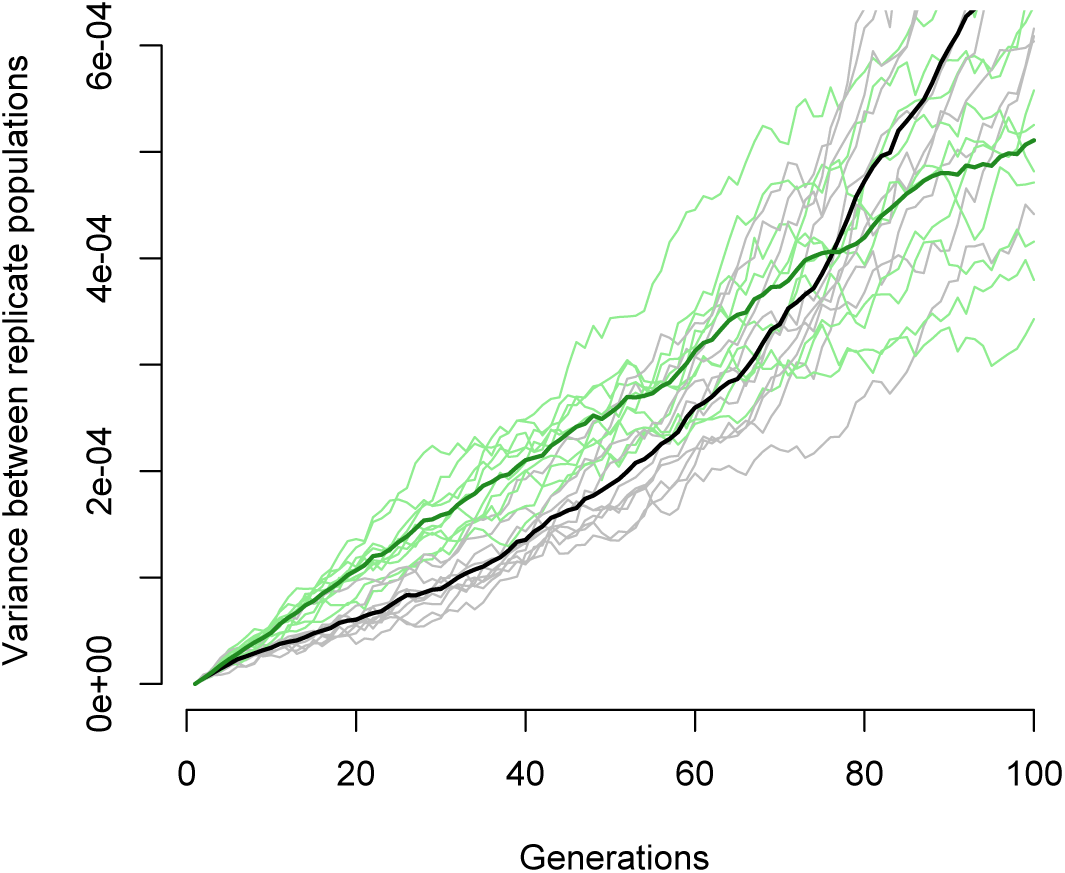
Expected differentiation in locomotion bias under an Ornstein–Uhlenbeck process. Vertical axis shows the variance in mean frequency of still individuals (as in Figure 1) across 50 replicates of the OU process in the selection surface of our experiment, assuming an effective population size of 1000 (Chelo and Teotónio, 2013) and the stable **G**-matrix of the domesticated A6140 population (Figure 3). Gray lines indicate results with selection and drift, green lines drift only. Thick black and green lines represent the average across 10 iterations of the simulations. Differentiation under drift is expected to follow a linear increase with time, while with selection it is better approximated by an exponential (Hansen and Martins, 1996). Results are similar when **G**-matrices other than the one from A6140 are used (not shown).

It is difficult to understand why phenotypic stasis is observed with the occurrence of both stabilizing and disruptive selection on phenotypic dimensions that are a complex combination of transition rates. While the addition of unobserved traits could change the genetic covariances between the observed traits, this is not so for the selection surface of locomotion bias in our lab environment; see for example discussion in Chapter 30 of Walsh and Lynch (2018). That is, the empirical limitations in characterizing the selection surface are only a function of the phenotypic space that we can measure with the inbred lines that were sampled. We suspect that selection must be strongly directional outside the small phenotypic space that we explored. We infer that populations meander in the phenotypic dimension under disruptive selection, mostly because of drift, but quickly slide off the slope once they approach a narrow optimum in the dimension of stabilizing selection. The speed at which populations showed stasis, soon after the hybridization of isolates, suggests this selection surface. Further sampling of wild isolates (Cook et al., 2017), and perhaps repeating experimental evolution with a more diverse hybrid population, would help address this hypothesis by exploring the phenotypic space beyond that we have mapped.

## 4 Summary and Conclusions

*C. elegans* isolates show great diversity in locomotion bias but it is apparent that soon after their hybridization phenotypic stasis occurred for 240 generations because of quadratic selection, presumably in concert with directional selection. On the other hand, the **G**-matrix of locomotion bias diverges from the ancestral state and differentiates among replicate populations facing the same constant laboratory environment within a period of 50 to 100 generations. During this period, mutation cannot account for divergence and differentiation of **G**. One of the central results of our study is that, instead, the evolution of the **G**-matrix is caused by genetic drift and quadratic selection, despite stasis at the level of individual trait means.

The specific problem posed by our findings is to understand how in the short-term of tens of generations, selection and drift generate enough linkage disequilibrium between QTL alleles such that genetic covariances between traits is not replicated among populations (Figure 5), without necessarily changing their trait means. Unfortunately, it is not immediately clear how the population genetics of QTL alleles translate into the maintenance of genetic variances and covariances (Kelly, 2009). Since QTL mapping models are undetermined, as the number of trait measurements were smaller than the genotypes that were assayed, and as the segregating variants at a particular time point of sampling cannot fully account for the distributions of the breeding values of previous generations, one must rely on simplified statistical descriptions of trait inheritance. Some of the larger effect QTL alleles still segregating after lab domestication are probably at low frequencies (Noble et al., 2019) but the implicit infinitesimal assumption of trait inheritance in our analysis might not hold in that the additive genetic variances and co-variances in locomotion will not be independent of allele frequency changes at smaller effect QTL. Genetic (co)variances are expected to be independent of allele frequency changes only in the long-term (i.e., thousands of generations, on the same order of the effective population size), when approaching mutation-selection-drift equilibrium (Vladar and Barton, 2014; Barton et al., 2017).

Assuming that after domestication a few QTL of moderate effects segregate at relatively low frequency, it is possible that selection and drift was such that each replicate population during divergence and differentiation had a greater chance of randomly fixing different alleles with pleiotropic effects that would not average out when comparing across populations. This would be akin to the process described by Cohan (1984). In the infinitesimal limit, an inflation of the effects of drift is indeed expected with selection on heritable quantitative traits because there is a correlation across generations between the breeding values of successful parents that results in a reduction in effective population sizes (Robertson, 1961). However, the Robertson (1961) model presupposes sequential, complete sweeps due to directional selection on initially rare QTL alleles, which may not be case when alleles are maintained at intermediate frequencies under stabilizing selection, as recently suggested with *Drosophila* experiments (Barghi et al., 2019) and likely to be the case here (Chelo and Teotónio, 2013). It is also possible, and perhaps probable in our setting, that small allele frequency and LD differences between replicate populations due to sampling of the domesticated population were accentuated by selection during the focal stage, resulting in variation between replicate populations of QTL effects on trait covariances (Bohren et al., 1966; Gomez et al., 2019). This process should be dampened by linked selection (Hill and Robertson, 1966; Zhang and Hill, 2005), although strong oligogenic sign epistasis we observed previously for fitness-related traits may exacerbate such sampling effects (Noble et al., 2017).

A better understanding of short-term phenotypic stasis with genetic divergence will come when the developmental biology of locomotion bias is better known (De Bono and Maricq, 2005; Zhen and Samuel, 2015), and explicit models of genotype to phenotype maps can be employed to predict phenotypic means from the genetic covariances of component traits (Riska, 1989; Rice, 2008). These approaches do not usually consider the interaction between genetic drift and selection on multiple traits. Rice (1998) suggested such modelling for “developmental systems drift” – the pattern that similar morphological phenotypes between species are realized through several redundant and potentially incompatible networks of gene expression (True and Haag, 2001; Johnson and Porter, 2007) – where interactions between developmental traits are neutral in some parts of the phenotypic space.

Modelling changes of allele effects with selection (Turelli, 1988; Shaw et al., 1995) likewise could be better understood through explicit models of development. In part, this is because in some of these models allelic effects can be scaled to population means (Hansen and Wagner, 2001; Morrissey, 2015). An example is given by Draghi and Whitlock (2012), who studied the evolution of phenotypic plasticity in changing environments to find that not only is the **G**-matrix poorly predictive of phenotypic responses to selection, but that only in the presence of genes that sense environmental cues will variation in the interactions between development traits result in the alignment of the **G**-matrix with the selection surface. This is a result expected from computer simulations of the adaptive landscape without explicit geno-type to phenotype maps, but only in the longer term when mutation starts to be significant (Jones et al., 2004, 2007).

We have shown that phenotypic stasis can occur with simultaneous divergence of the genetic variance-covariance structure of multiple component traits, as a product of quadratic selection and drift, and without limitation imposed by standing genetic variation. Phenotypic stasis is a common pattern observed in the evolutionary record over periods of up to 10^6^ years. For longer periods, rapid divergence in mean trait values is observed from fossil records, or inferred from phylogenetic trees, presumably because of sudden or extreme environmental change [see review in Arnold (2014)]. Our results suggest instead that upon environmental change, not necessarily sudden or extreme, populations might rapidly diverge and differentiate from each other due to past evolution of genetic variance-covariance structure. In addition, if standing genetic variation is lost under selection and drift, particular populations might be at risk of future maladaptation or extinction if the appropriate combination of component trait values is not expressed. If not at risk, then the pleiotropic properties of mutation should ultimately direct phenotypic evolution mostly independently of selection.

## 5 Materials and Methods

### 5.1 Experimental evolution

We analyzed the evolution of locomotion bias in a 240 generation experiment, the first 190 of which have been previously described (Teotónio et al., 2012; Theologidis et al., 2014; Noble et al., 2017). Briefly, 16 founders were inter-crossed to obtain a hybrid population (named A0), from which six population replicates (A[1-6]) were domesticated for 100 generations (140 generations for A[4-6] populations). From population A6 at generation 140 (A6140) we derived 5 replicate populations and maintained them in the same environment for another 100 generations (CA[1-5]). Recombinant inbred lines were generated by selfing from A6140, and from CA populations 1-3 at generation 50 and 100 (CA[1-3]50 and CA[1-3]100; Noble et al. (2019)). We refer to these 100 generations as the focal stage. During the domestication and focal stages, populations were cultured at constant census sizes of *N* = 10^4^ and expected average effective population sizes of around *N_e_* = 10^3^ (Chelo and Teotónio, 2013), under 4 day non-overlapping life-cycles (Teotónio et al., 2012), with periodic storage of samples (*>* 10^3^ individuals) by freezing (Stiernagle, 1999).

To estimate mutational effects on locomotion bias we assayed 250 generation mutation accumulation lines in two founder backgrounds (N2 and PB306), described in Baer et al. (2005) and Yeh et al. (2017).

### 5.2 Locomotion bias

#### 5.2.1 Assay block design

Hermaphrodite inbred lines from the experimental populations were phenotyped over 7 years in two different lab locations (Lisbon and Paris) by three different experimenters. All populations, the 16 founders, and mutation accumulation lines were phenotyped at a single location (Paris) by one experimenter (FM). We detail how covariate effects on locomotion bias were controlled in Section 5.2.6.

Samples were thawed from frozen stocks on 9 cm Petri dishes and grown until exhaustion of food (*Escherichia coli* HT115). This occurred 2-3 generations after thawing, after which individuals were washed from plates in M9 buffer. Adults were removed by centrifugation, and three plates per line were seeded with around 1000 larvae. Samples went through one to two complete generations in the controlled environment of experimental evolution, as detailed in Teotónio et al. (2012). At the assay generation (3-5 generations after thawing), adults were phenotyped for locomotion bias at their usual time of reproduction during experimental evolution (72h post L1 seeding) in a single 9 cm plate. At the beginning of each assay we measured ambient temperature and humidity in the imaging room to control for their effects on locomotion.

For the inbred lines derived from experimental populations there were 197 independent thaws, defining a statistical block, each containing between 2 to 22 samples (median=8). 189 inbred lines from the A6140 population were phenotyped, 52 from CA150, 52 CA250, 51 CA350, 51 CA1100, 53 CA1100 and 68 from CA1100. Each line was phenotyped in at least two blocks (plates), constituting a technical replicate (median=2, mean=2.25). For the experimental populations there were 9 independent thaws, of which 5 also contained founders. All founders and populations were phenotyped twice, with the exception of the ancestor population, A6140, which was included in 6 blocks. The CX12311 near isogenic line (kindly provided by P. McGrath) was phenotyped in a separate block along with the A6140 population for control. We phenotyped 58 and 66 mutation accumulation lines derived from the N2 and PB306 founders, respectively. All thaw blocks contained the N2 MA ancestor, and 15 of 16 blocks contained the PB306 ancestor. Full details of the number of observations for each genetic unit can be found in the archived tables (Section 9). Finally, and in order to improve the estimation of the individual fitness landscape in our experimental evolution environment (see below Section 5.4.3) we also assayed locomotion bias in 164 inbred lines derived from populations evolved in a high-salt environment (GA[1,2,4]50) for which fertility data was available for 56 (Noble et al., 2017).

#### 5.2.2 Worm tracking

To measure locomotion bias we imaged adults 72h post-L1 seeding using the Multi-Worm Tracker [MWT version 1.3.0; Swierczek et al. (2011)]. Movies were obtained with a Dalsa Falcon 4M30 CCD camera and National Instruments PCIe-1427 CameraLink card, imaging through a 0.13-0.16 mm cover glass placed in the plate lid, illuminated by a Schott A08926 backlight. Plates were imaged for approximately 20-25 minutes, with default MWT acquisition parameters: object size range of 50-6000 pixels, 10% object contrast, 50% fill hysteresis, 10% object size hysteresis, particle contours and skeletonization enabled. Choreography was used to filter and extract the number and persistence of tracked objects, and assign movement states across consecutive frames as forward, still or backwards, assuming that the dominant direction of movement in each track is forward (Swierczek et al., 2011).

MWT detects and loses objects over time as individual worms enter and leave the field of view or collide with each other. Each individual track is a period of continuous observation for a single object, so tracks that do not overlap in time might therefore correspond to the same individual worm. We ignored the first 5 minutes of recording, as worms are perturbed by plate handling (analysis not shown). Each movie contains around 1000 tracks with a mean duration of about 1 minute. Depending on the number of tracked objects (and computer performance), the MWT directly exports measurements at a frequency that can vary over time (Swierczek et al., 2011). Hence, in order to standardize the data we subsampled all movies to a frame rate of 4 Hz. Worm density, taken as the mean number of tracks recorded at each time point averaged over the total movie duration, was used as a covariate in the estimation of genetic variance-covariances (Section 5.3.2).

#### 5.2.3 Differentiating males from hermaphrodites

A6140 and all CA populations are androdioecious, with hermaphrodites and males segregating at intermediate frequencies (Teotónio et al., 2012; Theologidis et al., 2014). We were able to reliably (97% accuracy) differentiate between the sexes based on behavioural and morphological traits extracted from MWT data. In brief, we fit an initial three-trait Gaussian mixture model to control data, scored all samples by the difference in fit between one- and two-component models, then used a larger set of samples from the tails of this distribution to train an extreme gradient boosting classifier with R package xgboost (R Core Team, 2018; Chen and Guestrin, 2016) using a set of 30 derived traits.

In detail, we first evaluated a set of simple descriptions of individual size, shape and movement to find a subset of metrics that maximized the difference in preference for a two-component model between negative and positive controls: respectively, inbred wild isolate founders and two monoecious (M) populations which contained no, or very few, males; and three dioecious (D) populations with approximately 50% males (M and D populations were derived from A6140, see Theologidis et al. (2014) and Guzella et al. (2018)). Starting with worm area, length, width, curvature, velocity, acceleration, and movement run length as parent traits from Choreography output, child traits were defined by first splitting parents by individual movement state (forward, backward, still) and calculating the median and variance of the distribution for each track. Traits with more than 1% missing data were excluded, and values were log-transformed where strongly non-normal (a difference in Shapiro-Wilk −log10(p)>10). Fixed effects of block and log plate density were removed by linear regression before model fitting. Two-component Gaussian mixture models were fit to tracks for each line/population (R package mclust Scrucca et al. (2016), *V II* spherical model with varying volume), orienting labels by area (assuming males are always smaller than hermaphrodites). We sampled over sets of three traits (requiring three different parent trait classes, at least one related to size), and took the set maximizing the difference in median Integrated Complete-data Likelihood (ICL) between control groups (log area, log width, and velocity, all in the forward state). By this ranking, the 16 inbred founders and two monoecious populations fell within the lower 19 samples (of 77), while the three dioecious populations fell within the top 15 samples (although estimates appear to saturate at around 45% due presumably to density-dependent aggregation of males attempting to copulate).

To build a more sensitive classifier robust to male variation beyond the range seen in control data, we then trained an extreme gradient boosting model using the full set of 30 traits on the top/bottom 20 samples ranked by ICL in the three-trait mixture model [R package xgboost, Chen and Guestrin (2016)]. Negative control samples were assumed to be 100% hermaphrodite, while tracks in positive controls were assigned based on mclust model prediction (excluding those with classification uncertainty in the top decile). Tracks were classified by logistic regression, weighting samples inversely by size, with the best cross-validated model achieving an area under the precision-recall curve of 99.75% and a test classification error of 3.1% (*max*_*depth* = 4*, eta* = 0.3, *subsample* = 0.8*, eval*_*metric* = “*error*”). Prediction probabilities were discretized at 0.5.

Males tend to move much faster than females/hermaphrodites (Lipton et al., 2004), and since individual collision leads to loss of tracking sex is strongly confounded with track length and number. To estimate male frequencies at the sample level, tracks were sampled at 1s slices every 30s over each movie in the interval 400-1200 seconds, and line/population estimates were obtained from a binomial generalized linear model (Venables and Ripley, 2002).

#### 5.2.4 Modelling transition rates between locomotion states

We modelled the expected sex-specific transition rates between forward, still and backward states with a continuous time Markov process. Figure S2 shows the pipeline from the acquisition of movies to the final processed transition rate estimates.

We consider a system having *d* = 3 states with *P* (*t*_1_*, t*_2_)∈ℛ*^d,d^*, *t*_2_ *> t*_1_, denoting the transition probability matrix (Kalbfleisch and Lawless, 1985; Jackson, 2007):

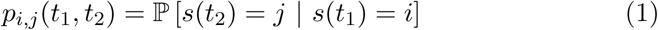

where *s*(*t*) ∈ ***S***, with ***S*** = {*<u>s</u>till, <u>f</u>orward, <u>b</u>ackward*} being the movement state occupied in instant *t*. We consider a time-homogeneous process described by the transition rate matrix:

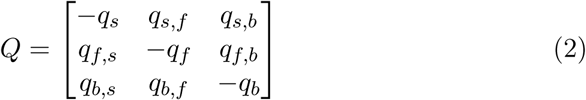

where *q_i,j_* ≥ 0 ∀*i, j*, subject to the constraint:

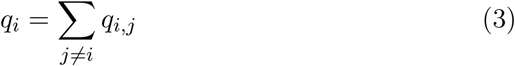

Hence, six of the nine possible transitions are independent. Let *θ* denote the parameters to be estimated, containing the off-diagonal elements from equation 2; this corresponds to:

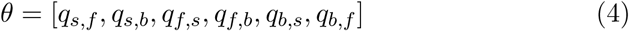

In this model, the time an object remains in a given state is on average 1*/q_i_*. Since the process is stationary, the probability of transition is a function of the time difference Δ*t* = *t*_2_ – *t*_1_, such that *P* (*t*_1_*, t*_2_) = *P* (Δ*t*), and the elements of the *P* (Δ*t*) matrix are given by:

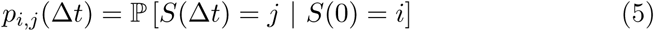

It then follows that:

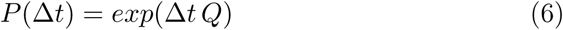

where exp(.) denotes the matrix exponential. The constraint in equation 3 ensures that:

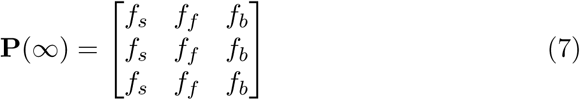

where *f_i_* is the relative frequency of state *i* that no longer depends on the previous state (all three rows of the **P**(∞) matrix converged). We find that the state frequencies from **P**(∞) are a monotonic and mostly linear function of the observed frequencies of movement states (Figure S3), showing that violations of the Markov assumption of the model does not induce a large bias in the long-term predictions of our model.

#### 5.2.5 Estimating transition rates

In our case, we have *N* objects (individual tracks; see above Section 5.2.2) for a single plate (technical replicate), with the data on the *k*-th object denoted as:

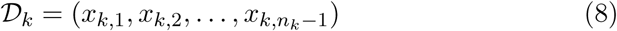

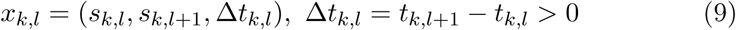

where *s_k,l_* is the state of the *k*-th object in the *l*-th time-point in which it was observed, and *t_k,l_* is the instant of time in which this observation was made. Then, given data 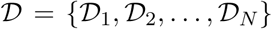, the log-likelihood for the model is given by (Kalbfleisch and Lawless, 1985; Bladt and Sorensen, 2005):

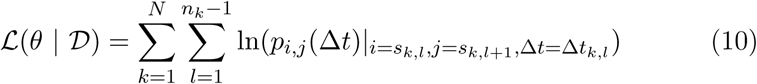

where *p_i,j_* (Δ*t*) was defined in equation 5, being calculated as a function of the parameters *θ* via equation 6. Therefore, the data on the *N* objects can be represented as the number of observations of *x* = (*i, j,* Δ*t*), which we denote as *ñ_i,j,_*_Δ_*_t_*:

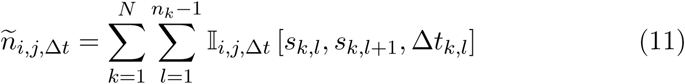

where I*_i,j,_*_Δ_*_t_* [·] is the indicator function:

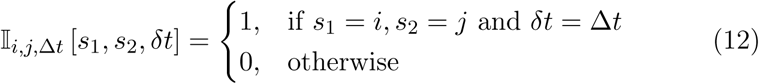

Under this model the input data can be compressed by considering only the data:

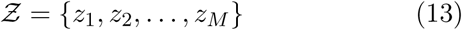

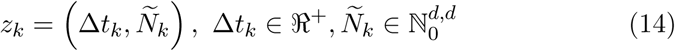

where:

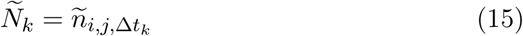

Finally, the log-likelihood can be rewritten as:

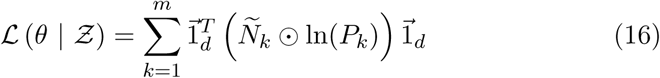

where 1*_d_* is a *d*-dimensional vector of 1s, denotes ʘ the Hadamard prod uct, and ln *P_k_* is the matrix obtained by taking the logarithm of each value in matrix *P_k_*.

Models were specified using RStan (Stan Development Team (2018), R version 3.3.2, RStan version 2.15.1), which performs Bayesian inference using a Hamiltonian Monte Carlo sampling to calculate the posterior probability of the parameters given the observed data. We used multi-lognormal prior distibutions with mean transition rate of 2 s*^−^*^1^ and a coefficient of variation of 60%:

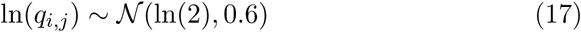

Throughout the rest of the methods section, we will denote *q_k_* the six off-diagonal elements of the **Q** matrix estimated by this model.

Three separate iterations of the model were run containing the data from (1) inbred lines (2) populations and founders and (3) MA lines. This does not affect the final results as the transition rates for each plate are independent estimates in the model. Final plate-level estimates and all subsequent R code are available from our github repository (Section 9).

#### 5.2.6 Controlling for experimenter and time of data acquisition

With per-plate estimates for transition rates, we fitted preliminary models to account for the effects of assay design and homogenize data sets. We first corrected for experimenter effects using a set of 26 inbred lines that were measured by all three experimenters. For each transition rate we fitted the linear mixed-effects model (*lmer* function in R package *lme*4 Bates et al. (2015)):

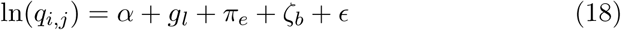

with *α* being the mean trait value of all the lines, *g* fixed effects of each line *l*, *π* fixed effects of experimenter *e*, *ζ ∼ N*(0*, σ*^2^) a random effects of block *b*, and *ϵ* ∼ *N* (0*,σ*^2^) the residual error. The same procedure was used to correct for a significant location effect, using a shared set of 64 inbred lines:

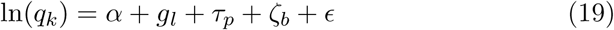

with *τ* the fixed effect of location *p*. The estimated experimenter and location effects from these models were then used to adjust all data for subsequent analysis.

#### 5.2.7 Repeatability

Transition rate repeatabilities were computed from linear mixed models using the R package *rptR* (Stoffel et al. (2017)), with confidence intervals estimated from 1000 bootstrap samples. We used the univariate versions of the models presented in equation 21 and 23 to estimate the repeatability of A6140, CA[1-3]50 and CA[1-3]100 RILs and populations, and the MA lines from each ancestor (N2 and PB306), presented in Figure S15.

### 5.3 Genetic variance-covariances of transition rates

The inbred lines derived from focal experimental populations allowed estimation of the **G**-matrix for the six non-self transition rates *q_i,j_* as the random genetic differences between lines after accounting for covariates (Lynch and Walsh, 1998). The diagonal entries of **G** are the (additive) genetic variance for each transition rate, while the off-diagonal entries are the (additive) genetic covariances between transitions rates. Likewise, the mutation accumulation lines allow us to calculate the **M**-matrix, giving the expected random genetic variances and covariances of transition rates due to mostly unselected mutations fixed after 250 generations of evolution at an effective population size close to 1 (Baer et al., 2005).

#### 5.3.1 Inbreeding effects

In the presence of directional dominance and epistasis, the additive genetic (co)variances estimated with the inbred lines are only an upper bound on the true values of the respective outbred populations (Lynch and Walsh, 1998). To gauge the extent of inbreeding effects on transition rates we compared the mean values of experimental populations with those of the inbred lines derived from them. For the populations, confidence intervals were extracted from the mixed model using the R function *confint*, for the inbred lines intervals, assuming normality, are the mean ±1.96×*σ*^2^. Significant differences are indicative of directional dominance and/or epistasis, as genotypes expressing non-additive effects are fixed with inbreeding (Lynch and Walsh, 1998).

#### 5.3.2 G-matrix estimation

Statistical analysis used the R package MCMCglmm (version 2.26, MCM-Cglmm function with default parameters; Hadfield (2010)). We first removed the effects of environmental covariates recorded during data acquisition (Section 5.2.1) for each of three groups of populations (A6140, CA[1-3]50, CA[1-3]100), using the generalized linear mixed-effects model for each population and trait:

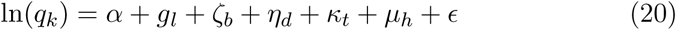

with *g* and *ζ* as defined above (Section 5.2.6), and *η*, *κ* and *µ* the fixed effects of ln(density) *d*, temperature *t* and relative humidity *h* respectively (Sections 5.2.1 and 5.2.2). **G**-matrices were then estimated separately for each of the seven experimental populations (A6140, CA[1-3]50, CA[1-3]100). With **Y** the *q* × *l* matrix of *q*(= 6) transition rates and *l* inbred lines:

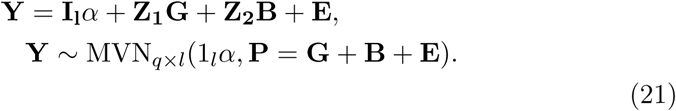

with **I** the identity matrix, *α* the matrix of mean trait values for each line, **Z**_1_ the design matrix of random genetic effects (line identities) and **Z**_2_ the design matrix for random block effects. In this model, the phenotypic (co)variance matrix **P** is decomposed into genetic (**G**), block (**B**) and residual (**E**; comprising variance due to uncontrolled effects of environment, non-additivity and measurement error) components.

We constructed priors as the matrix of phenotypic variances for each trait, ignoring covariances as recommended by Wilson et al. (2010). Model convergence was verified by visual inspection of the posterior distributions and by ensuring that autocorrelation remained below 0.05. We used 50,000 burn-in iterations, then took samples every 10 iterations over the next 100,000.

#### 5.3.3 M-matrix estimation

The MA lines were phenotyped in separate blocks. For each transition rate we first modelled block effects (*ζ*) using the data obtained from the two ancestral lines (n=31) in a linear model:

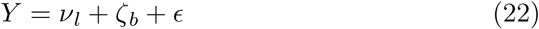

with *ν* the mean value of each ancestral line *l* (N2 or PB306). Block effects were then used to correct the values of all MA lines before computing the **M**-matrices. We fitted fixed effects of environmental covariates (density, temperature and relative humidity) within each ancestral sub-group of mutation accumulation lines in the multivariate mixed model as for **G**-matrices:

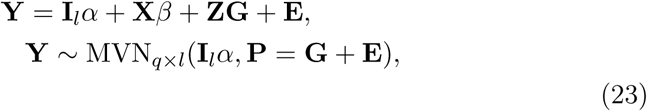

with the addition of the design matrix **X** for fixed covariate effects in *β* (comprising, as above, *β_d_*, *β_t_* and *β_rh_*) for all traits. **Z** is the design matrix of random genetic effects (line identities).

#### 5.3.4 G-matrix divergence

There is a large variety of approaches to test for differences between **G**-matrices (Aguirre et al., 2014). We present results for two: the pairwise Flury hierarchy (Phillips and Arnold, 1999), and a joint tensor analysis (see Appendix 6).

The Flury comparison consists of a nested succession of hypothesis tests concerning the degree of similitude between two matrices. In this hierarchy (from top to bottom), two matrices are considered equal, proportional, similar in sharing some number of common principal components (CPC) or, finally, unrelated. We used CPCRAND to perform these tests (Phillips, 1998), which applies the randomization methods described in Phillips and Arnold (1999) while applying bootstrapping for error estimation. Descending through the hierarchy, each hypothesis is tested against the null hypothesis at the bottom of the hierarchy (i.e., unrelated), continuing until the hierarchy is traversed, or the null is rejected.

We calculated the orthogonal axes of **G**-matrices (base R function *eigen*) in which genetic variance is maximized (Schluter, 1996), and that indicate how **G** is expected to influence the speed and dynamic of responses to selection. We represent in Figures 4 and 6 the angle between eigenvectors calculated as:

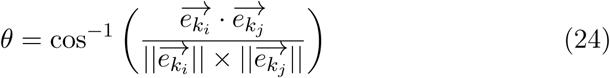

with 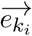 and 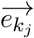 being the *k*-th eigenvectors of the two matrices to compare.

#### 5.3.5 Expected G-matrix evolution by genetic drift

We designed individual-based Monte Carlo simulations to estimate the change in the ancestral A6140 **G**-matrix expected with 100 generations of genetic drift (the time frame of the focal stage, Section 5.1). The model we employ is similar to that of Jones and colleagues, e.g. Jones et al. (2007, 2014), although our framework allows modelling of up to six traits and variable pleiotropy defined at the allele level. Simulations were coded in R (Section 9).

We simulated diploid individuals for a total population size of 1000 (the estimated effective population sizes of our populations from Chelo and Teotónio (2013)). Genomes were composed of 100 unlinked loci, each starting with 16 possible founder alleles, with each allele affecting one to six traits (sampled from a truncated Poisson distribution with mean=3. Varying this number does not affect the conclusions, not shown). Allelic effect sizes were randomly sampled from a multivariate normal with mean equal to zero.

We further conditioned starting allelic effects on **G**-matrices similar to that of the experimental ancestor A6140. Only with maximal pleiotropy, where all alleles affect all traits, would we be able to recreate the observed A6140 **G**-matrix. We instead scaled allelic effect sizes (i) in proportion to the number of traits they affect, (ii) inversely proportional to their expected frequency, and (iii) by a global scaling value applied to all effects, as determined in preliminary simulations minimizing the difference between simulated ancestral and observed A6140 **G**-matrices (not shown). The simulated ancestral allele frequency was based on the empirical SNP frequency distribution of the A6140 population estimated from inbred lines Noble et al. (2019).

This distribution is well-approximated by an exponential with mean=5, to which was added 0.15 to account for the most frequent allele. We then truncated this distribution by retaining values below 0.9 in order to ensure that all sites have segregating alleles in the ancestral populations. Using this approach, for each locus we randomly attributed each allele to the individual of the ancestral population for simulations.

Once the ancestral population for the simulations was created, we iterated outcrossing 100 times: (i) individuals mate, (ii) each mated pair produces exactly 10 offspring (although in excess of the per-pair expected contribution of 2 offspring, lack of variation between pairs will diminish the effects of drift), and (iii) 1000 individuals are retained from the progeny pool, corresponding to the estimated effective population size during experimental evolution (Chelo and Teotónio, 2013).

To compare observed and simulated **G**-matrices we replicated the assay sampling scheme, randomly selecting 100 simulated chromosomes representing fully inbred individuals, and then summing allelic effects for each individual in the natural linear scale. Resulting **G**-matrices were modelled as above using a multivariate normal linear mixed-effects model (Section 5.3.2).

#### 5.3.6 Modelling G-matrices with genomic data

We have previously described genotyping by genome sequencing of the inbred lines phenotyped for locomotion bias (Noble et al., 2019). To estimate genomic **G**-matrices from realized genetic relatedness we fit multivariate linear mixed-effects models for each experimental population [R package sommer, Covarrubias-Pazaran (2016)], using an additive genetic relationship matrix **A** generated from the SNPs of Noble et al. (2019) (data set 1), and trait values adjusted for fixed effect covariates (Methods 5.2.6). The univariate model fit for trait 1 (of *p* = 6) is:

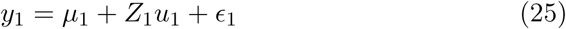

where *µ* is the trait mean (intercept), *Z* is a design matrix for random effects, *u* and *ϵ* are *n* random genetic and residual effects, respectively, assumed independently and normally distributed with zero mean. Following (Maier et al., 2015), the phenotypic variance-covariance matrix, comprising additive genetic and residual matrices (in **G** and **E**), is:

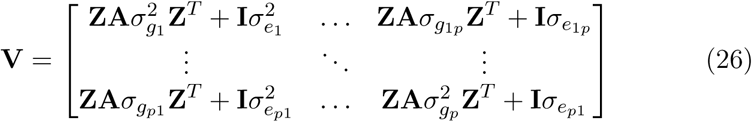

where 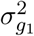 and *σ_g_*_1_*_p_* are genetic variances for trait 1 and genetic covariances between traits 1 and *p*, respectively (with corresponding residual values). With **S** the *m* marker by *n* line matrix of centered and scaled geno-types, **A=SS^T^***/m*. Residual maximum likelihood (REML) estimates were obtained by the Newton-Raphson algorithm.

To test for effects of variation in LD between populations on **G**, **A** matrices were constructed for each population by independently pruning segregating markers of pairwise linkage disequilibrium within each chromosome at increasing maximum *r*^2^ thresholds (*window* = 1500 × *r*^2^*, step* = 1000 *r*^2^). This was repeated 10 times for each **A** randomly retaining markers below the threshold in each window each run. Figure 5A summarises mean pairwise differences in these **G**-matrices by normalising first to each value obtained when using all markers to construct **A**, for each population, then subtracting the value for the ancestral population.

### 5.4 Selection on locomotion bias

#### 5.4.1 Retrospective analysis

We first computed the average value for each founder, replicate population, and time point by fitting a linear mixed-effects model using the *lmer* function in R:

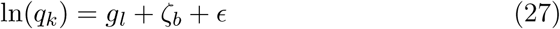

with the factors being defined as in equation 20, *g* being the estimate for each population or wild isolate *l* (n=72). We did not include environmental covariates because all measurements were done in a short period of time and by a single experimenter (FM) so that the environmental variation is captured by the among-block variation (n=10 out of 197 in total). The same model was then used to compute the mean population value of the mean frequency of the three movement states (still, forward and backward). Males and hermaphrodites were analyzed separately.

Following Lande (1979), we estimate net gradients of directional selection (*β_net_*) on each transition rate as:

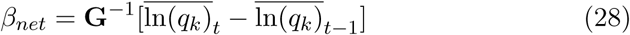

where 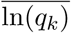 is the mean trait value of a given replicate population, compared at generation *t* and the previous time point *t –* 1 (or the ancestral A0 or A6140 population; from equation 27). We sampled variable numbers of replicate populations at different generations for a total of 49 estimates for each transition rate (Figure 1). This approximation for directional selection holds if the **G**-matrix is constant during the period considered (Turelli, 1988; Shaw et al., 1995) and, without loss of generality, in this analysis we considered that **G** was the identity matrix.

#### 5.4.2 Selection surface

In Noble et al. (2017) we reported hermaphrodite fertility data on 230 inbred lines derived from several populations. Fertility includes fecundity and offspring survival to the L1 larval stage, and was measured in environmental conditions that closely followed those of experimental evolution (except for interaction with males and final larval density). We have no prior indication that hermaphrodite fertility trades-off with viability (Chelo et al., 2019), and it therefore represents a major fitness component that we use here as a fitness surrogate.

The log-transformed, covariate-adjusted fertility values for each line were downloaded from Noble et al. (2017), exponentiated, and divided by the sum of all values to obtain a relative fitness measure (*w*). For the six transition rates, we first obtained the mean value for each line by fitting a linear mixed-effects model using the *lmer* function in R:

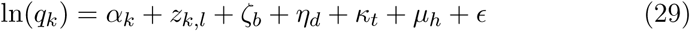

with the factors being defined as in equation 20, and estimates of the interaction between the covariates density, temperature and humidity being retrieved. Because *α_k_* are the overall mean trait values, the computed mean transition rates per line (*z_k,l_*) are centered.

To estimate the selection surface (also called the “individual fitness function” or the “fitness landscape”), we used partial regression of the relative fitness of each line (*w_l_*) on transition rates (*g_k,l_*):

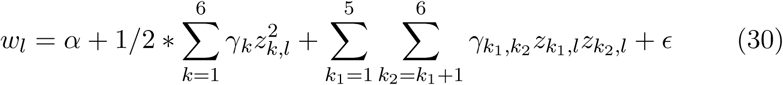

This model approximates the curvature and tilt of the six dimensional selection surface (Lande and Arnold, 1983), with *α* being the vector of mean transition rates among lines and *γ* the partial regression coefficients estimating quadratic and interaction (stabilizing or disruptive, depending on the sign) selection on each transition rate *k*, or correlated selection between pairs of transition rates *k*_1_ and *k*_2_. Estimates from a full model including linear coefficients are shown as supplementary information (Figure S10), since their inclusion did not greatly affect inferences.

The regression model was fitted with RStan (Section 9) with improper priors, and convergence was checked by visual inspection of the posterior distributions and by ensuring that autocorrelation remained below 0.05. Fitting the same model using maximum likelihood (*lm* function in R) gave similar results (not shown).

#### 5.4.3 Canonical analysis of the selection surface

The selection surface is given by the symmetric ***γ***-matrix, with *γ_k_* coefficients on the diagonals and *γ_k_*_1_*_,k_*_2_ coefficients on the off-diagonals. We used canonical analysis (eigen decomposition) (Phillips and Arnold, 1989) to characterize the main selection axes on (weighted) combinations of transition rates:

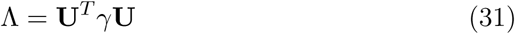

with **U** being the matrix of eigenvectors of ***γ***, and Λ the diagonal matrix of eigenvalues (denoted *λ*[_1_*_−_*_6_]). As for *γ*, the signs of the six *λ* coefficients define the curvature of the fitness landscape around equilibria, with positive (negative) values indicating fitness maxima (minima).

#### 5.4.4 Transition rate projections onto the selection surface

The **U**-matrix in equation 31 can be used to rotate the vector of mean transition rate values (*z_k_*) onto a new canonical trait vector (*y_k_*) along the eigenvectors of ***γ***:

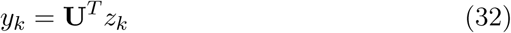

Under this transformation, the relative fitness e quation becomes:

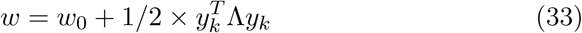

with *w*_0_ the fitness at the stationary point. To represent the phenotype distribution of genetic entities on the fitness landscape, we centered and rotated all measurements using equation 32; the model in equation 29 being applied beforehand to obtain population and MA line mean estimates. Similarly, genetic variance-covariance matrices were rotated to visualize them on the fitness landscape:

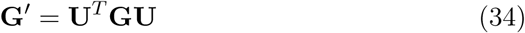

and

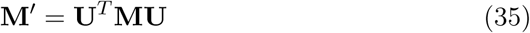

For Figures 8, S11 and S14, we estimated the amount of genetic variance maintained by selection by regressing the sum of the absolute values of the column entries in **G***^t^* or **M***^t^* (genetic variance-covariances of each *g′_i,j_* canonical trait) on the strength of selection |*λ*|, sampling the posterior distributions of the estimated **G**, **M** and ***γ*** matrices 100 times.

### 5.5 Modelling divergence of trait means

We modelled the evolution of mean population trait values as a Ornstein–Uhlenbeck (OU) process, which have been used to model phenotypic responses around an optimum (Felsenstein, 1988; Hansen and Martins, 1996). During this process, the mean trait values change over generations by Brownian motion, to which is added a restraining selective force that pulls the phenotype towards the equilibrium, or pushes them away from it. Under the assumption that selection is weak, selection strength is proportional to **G***γ* and increases as the trait mean diverges from the phenotypic optimum *θ*.

Moreover, assuming constant **G**, **P** (the phenotypic variance-covariance matrix), *γ*, *θ* and *N_e_*, then *X*(*t*), the vector of trait means at a given time, can be assumed to follow a OU process with infinitesimal mean **G***γ* and variance **G**/*N_e_*. In this case the vector of trait means at the next generation is:

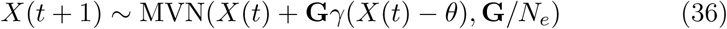

We applied this model to our data, using as the optimum *θ* the mean trait values used for centering during *γ* estimation, and the **G**-matrix of the A6140 population. Starting from the phenotypic optimum, we generated 50 independent time series of 100 generations at *N_e_* = 1000. For weak selection *β ≈* 0 and *γ <<* **P**, and while we have shown that *β ≈* 0, the eigenvalues of *γ* are larger than the phenotypic variance. The same OU process was then iterated assuming only genetic drift with the vector of trait means being:

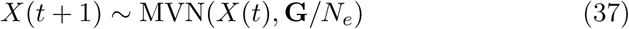

## 6 Appendix: tensor analysis of G-matrix divergence

Genetic covariance tensors (**Σ**) are fourth-order objects that describe the variation among multiple **G**-matrices (Hine et al., 2009; Aguirre et al., 2014). As for dimensional reduction of second-order matrices along principal components, the variation among multiple G-matrices captured in **Σ** can be described by decomposition into orthogonal eigentensors (**E**), where each **E** is associated with an eigenvalue that quantifies its contribution to variation in **Σ**. In turn, eigentensors **E** are second-order objects that can be decomposed into eigenvectors (e, each associated with an eigenvalue).

Aguirre and colleagues have implemented this approach in a Bayesian framework that accounts for uncertainty in **G**-matrix estimation (Aguirre et al., 2014). The amount of variance *α***_i_** associated with each eigentensor **E_i_** is compared to the expected *α***_i_** under a null model where the variation among matrices is due to sampling variation. In our data, we found that the first two eigentensors, **E_1_** and **E_2_**, explain more variation than expected under the null (Figure A1). We further tested (i) whether **G**-matrices diverge in the same directions or (ii) whether replicate populations change in different directions. In the former scenario, we should observe a difference between the ancestral **G**-matrix and those of the evolved populations, and this difference might scale with time. In the latter case, we should not observe any consistent change. We examined the contribution of each population to the eigenvalues of the tensor by plotting the coordinates of each **G** in the space of these two significant eigentensors (Figure A2): while there is considerable uncertainty in the estimates, no single population seems to drive most of the variation among the G-matrices. In the space of the eigentensors **E_1_** and **E_2_**, the ancestral population shows the highest absolute values in both spaces. After 50 generations evolutionary trajectories are inconsistent among replicates: CA250 and CA350 are the most divergent populations in the first space, while in the second space it is CA150. The same pattern is seen at 100 generations, where divergence from the ancestral population is inversely correlated across the two spaces.

Finally, we examined the contribution of specific trait combinations to coordinated change among **G**-matrices. We projected the eigenvectors of each significant eigentensor on the observed **G**-matrix, giving estimates of genetic variance in each population in the direction of the greatest variation among **G** (Figure A3). The first two eigenvectors of *E*_1_ and *E*_2_ are associated with a general reduction in variance from the ancestral population, which varies among replicates along different directions. For the first eigentensor **E_1_**, loss of variance is negatively correlated in the direction of *e*11 and *e*12. For example, after 50 generations of evolution we can see that CA250 mostly lost variance along *e*11 while CA150 did so along *e*12. Interestingly, *e*11 and *e*12 are highly correlated with *y*_3_ and *y*_2_, the second and third eigenvectors of our *γ*-matrix (*r* = −0.92 and 0.94, respectively). In the second eigentensor **E_2_**, most of the populations lost variance along *e*21 that correlates weakly with three eigenvectors of *γ* (*y*_2_,*y*_3_ and *y*_5_ with all 3 correlation coefficients *r ≈*0.6). The exception is CA350, that mostly lost variance along *e*22, strongly correlated with *y*_4_ (*r* = 0.93).

The tensor analysis shows that our **G**-matrices diverged during experimental evolution, but we do not find concordant changes across replicates or time. The eigentensor axes where most change is observed correspond to those of the selection surface where most of the variation in **G** is due to loss of genetic variance along specific trait axes under weak selection. The observation that replicate populations diverged in different phenotypic directions, and that these changes are not consistent in time, further suggests that drift has played the dominant role in this area of trait space.

**Figure A1:**
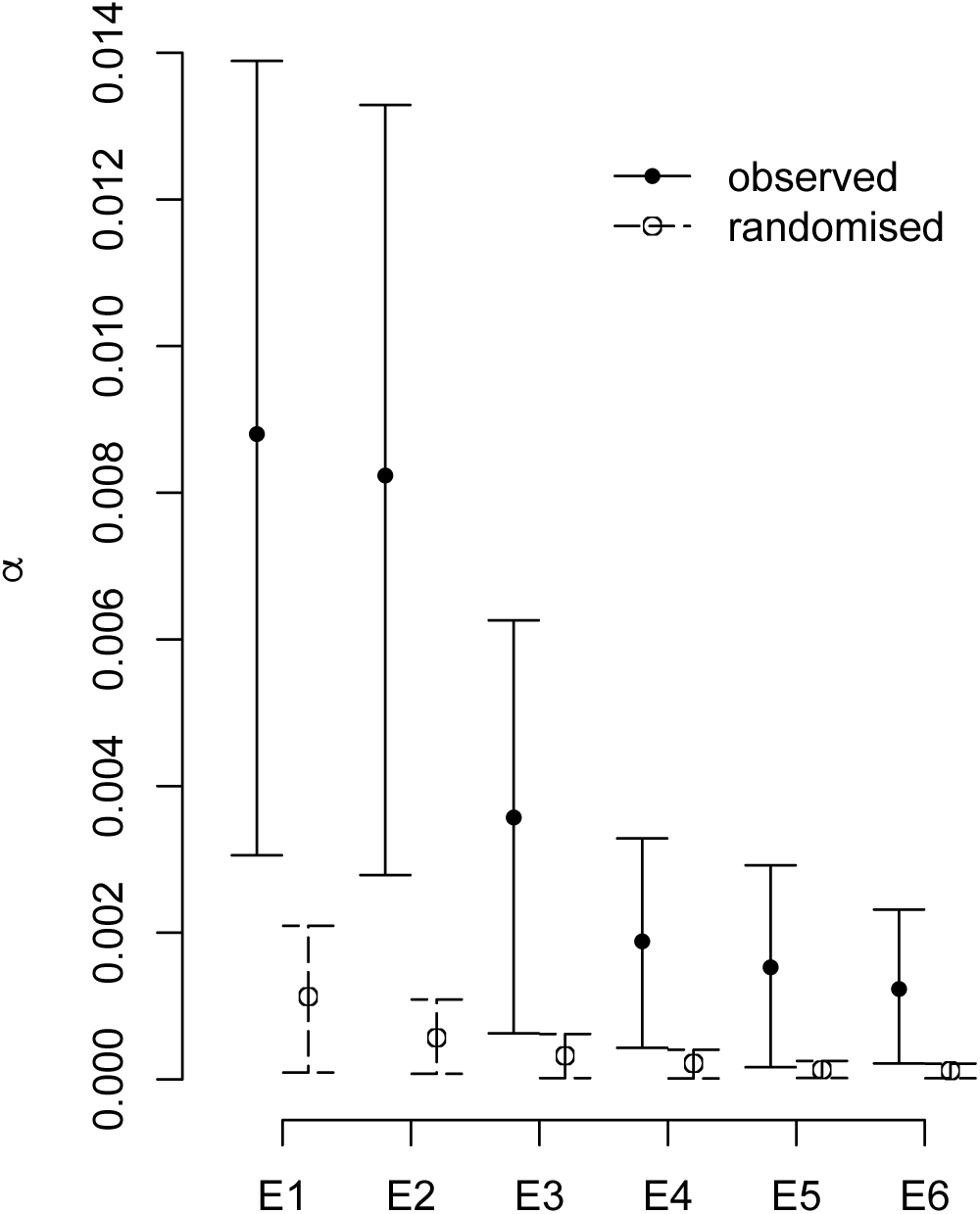
Variation among observed and null **G**-matrices along each eigentensor. The variance *α***_i_** associated with each eigentensor **E_i_** is compared to the expected *α***_i_** under a null permutation model where variation among matrices is due to sampling variation. Here, the first two eigentensors, **E_1_** and **E_2_**, show levels of variation that are unlikely under the null (bars show 90% confidence intervals).

**Figure A2:**
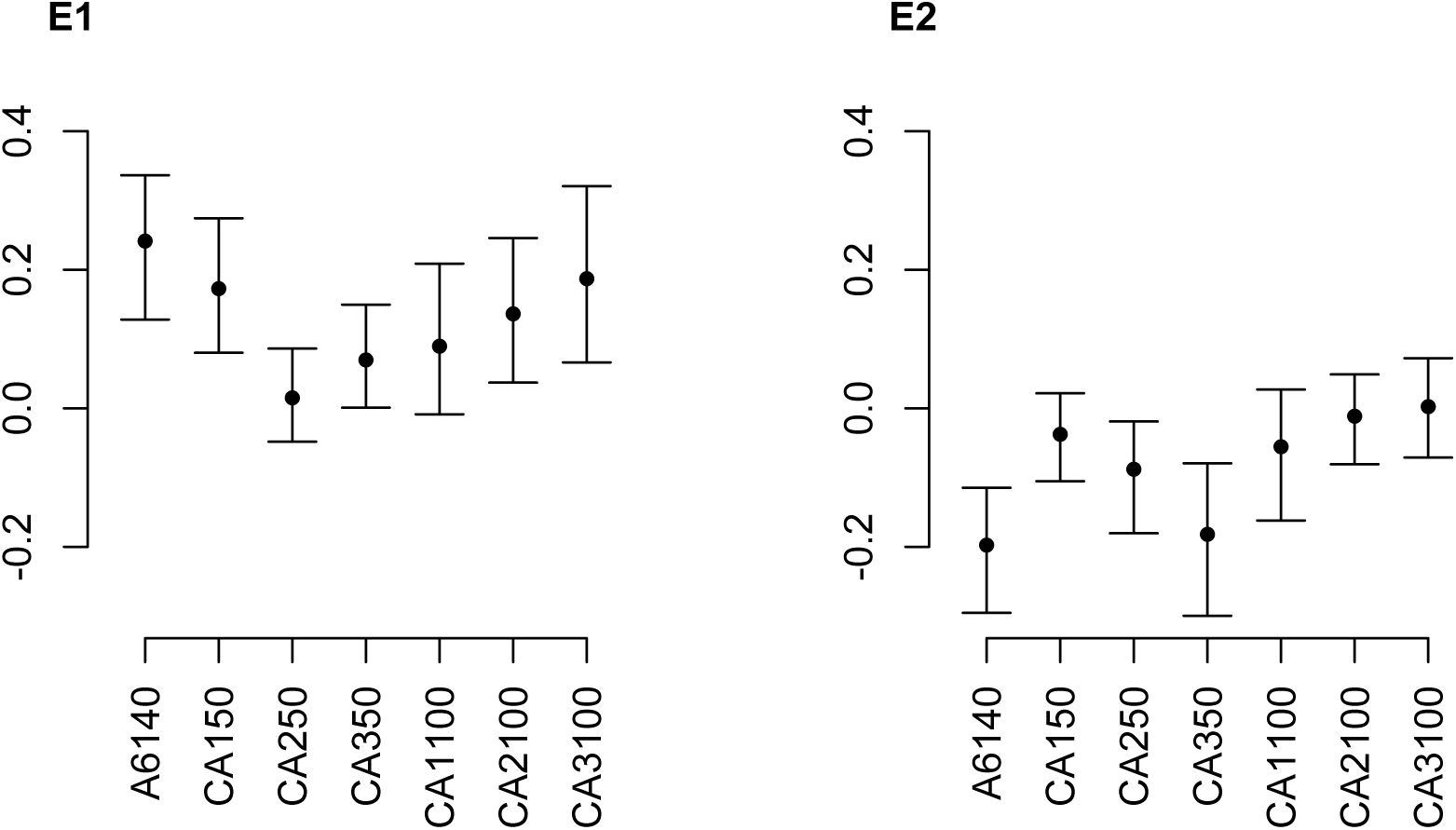
Coordinates of the posterior mean **G**-matrices and 95% highest posterior density (HPD) intervals of MCMC samples in the space of **E_1_** and **E_2_** for the posterior mean of each population. In both spaces, the ancestral population A6140 has the largest absolute value showing that it drives most of the variation seen in Figure A1. Within these spaces, the pattern of variance loss is uncorrelated with time or replicate lineage.

**Figure A3:**
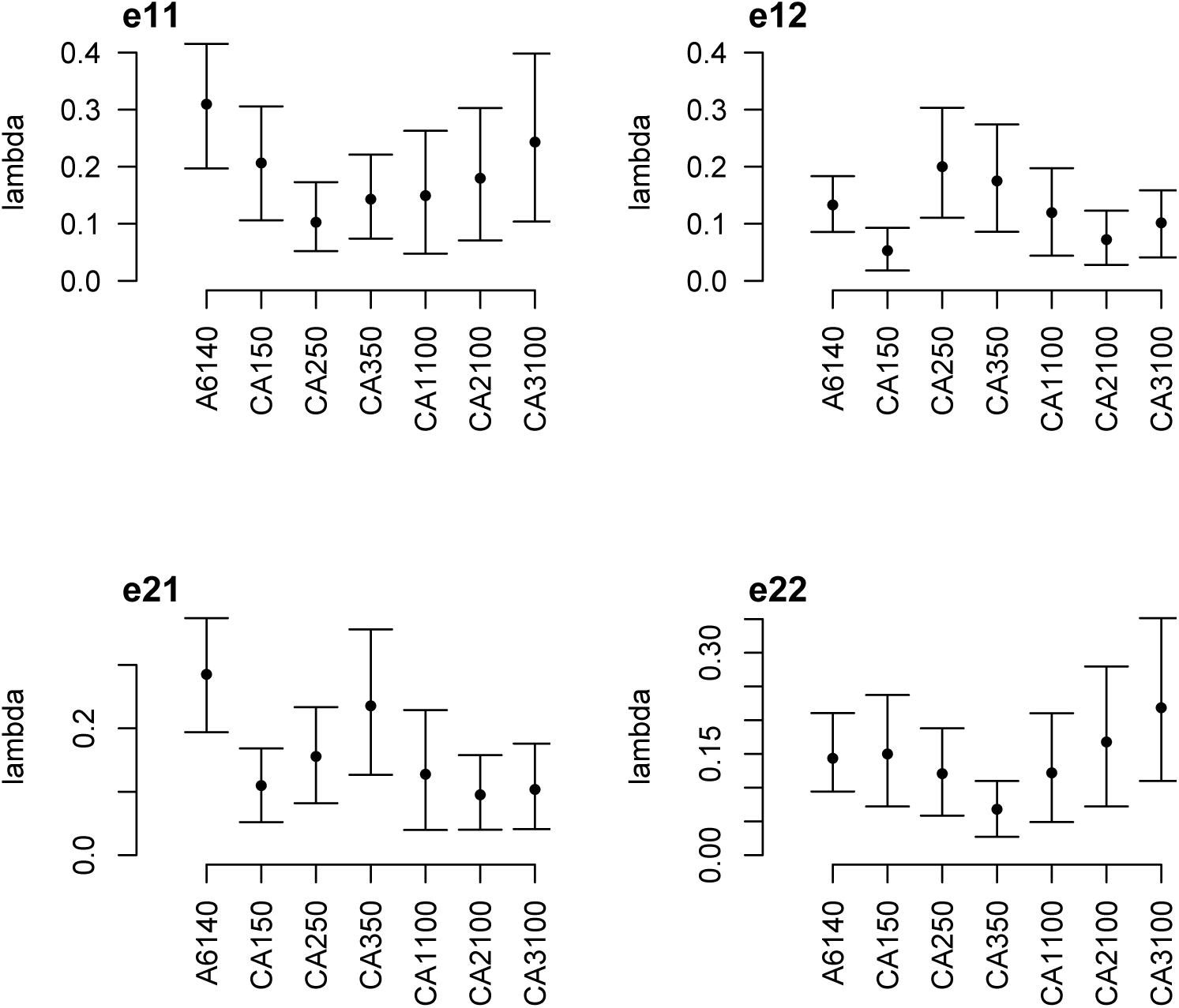
Contribution of specific trait combinations to coordinated changes among **G**-matrices. Each panel shows the amount genetic variance for each population in the direction of the greatest variation among **G** (eigenvectors of **E_1_** and **E_2_**).

## 7 Supplementary Figures

**Figure S1:**
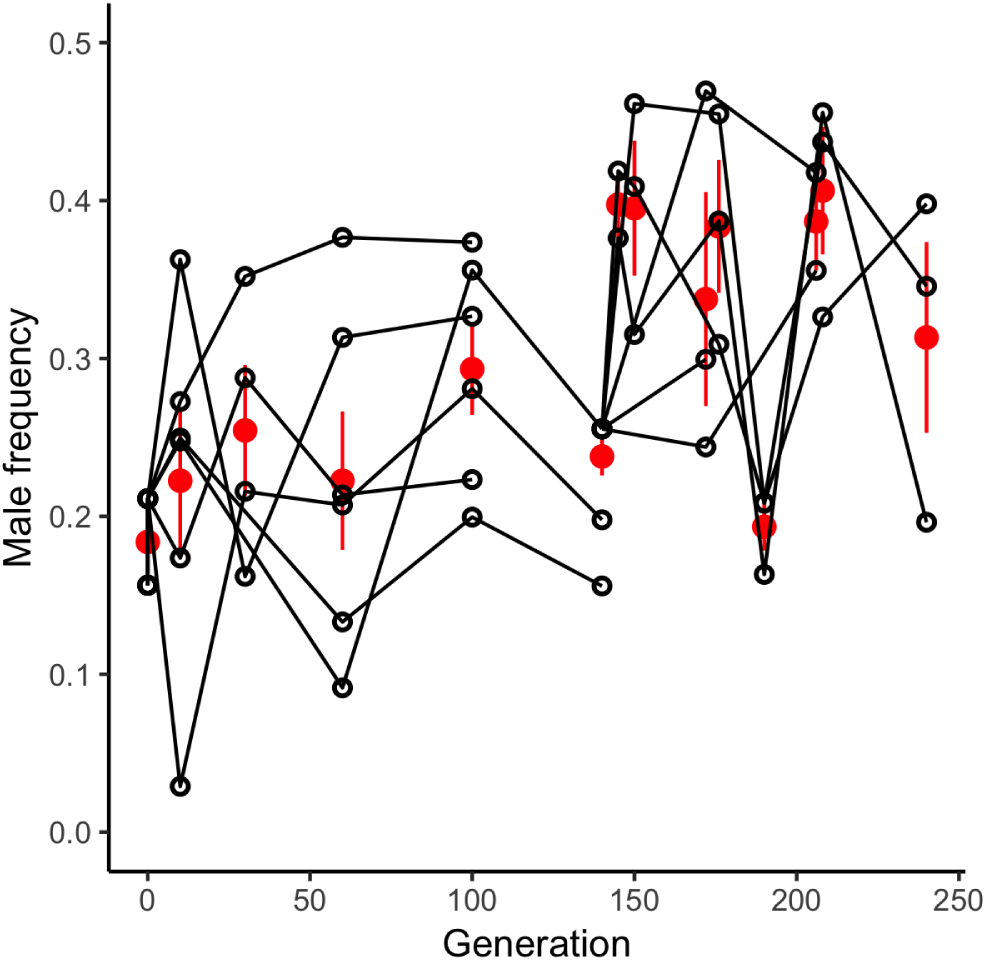
Male frequencies during experimental evolution. Males and hermaphrodite tracks were differentiated with a 30-trait classifier based on moments of size, shape and velocity-related traits derived from Multi-Worm Tracker metrics, and frequencies were estimated from 1s slices across movies (Methods 5.2.3). Empty circles indicate the estimates for each replicate population (between 1 and 6 at each time point), red circles the mean among replicate populations (±standard error). During the first 100 generations of domestication, the estimates are similar to those obtained by directly counting the number of males (Teotónio et al., 2012). As *C. elegans* is androdioecious, with hermaphrodites being able to self or mate with males, expected outcrossing rates are twice the male frequencies.

**Figure S2:**
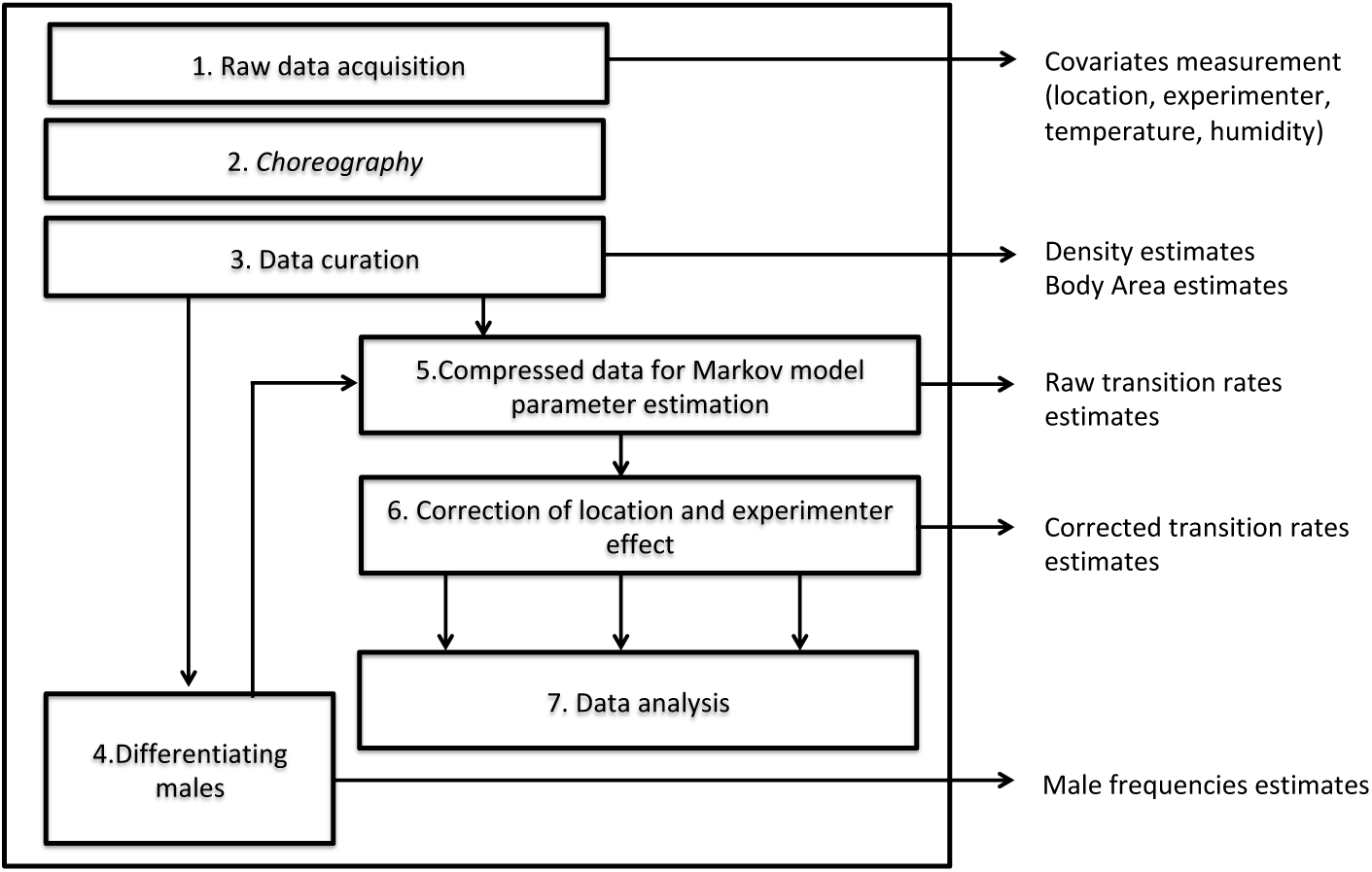
Schematic of phenotype data acquisition and analysis pipeline.

**Figure S3:**
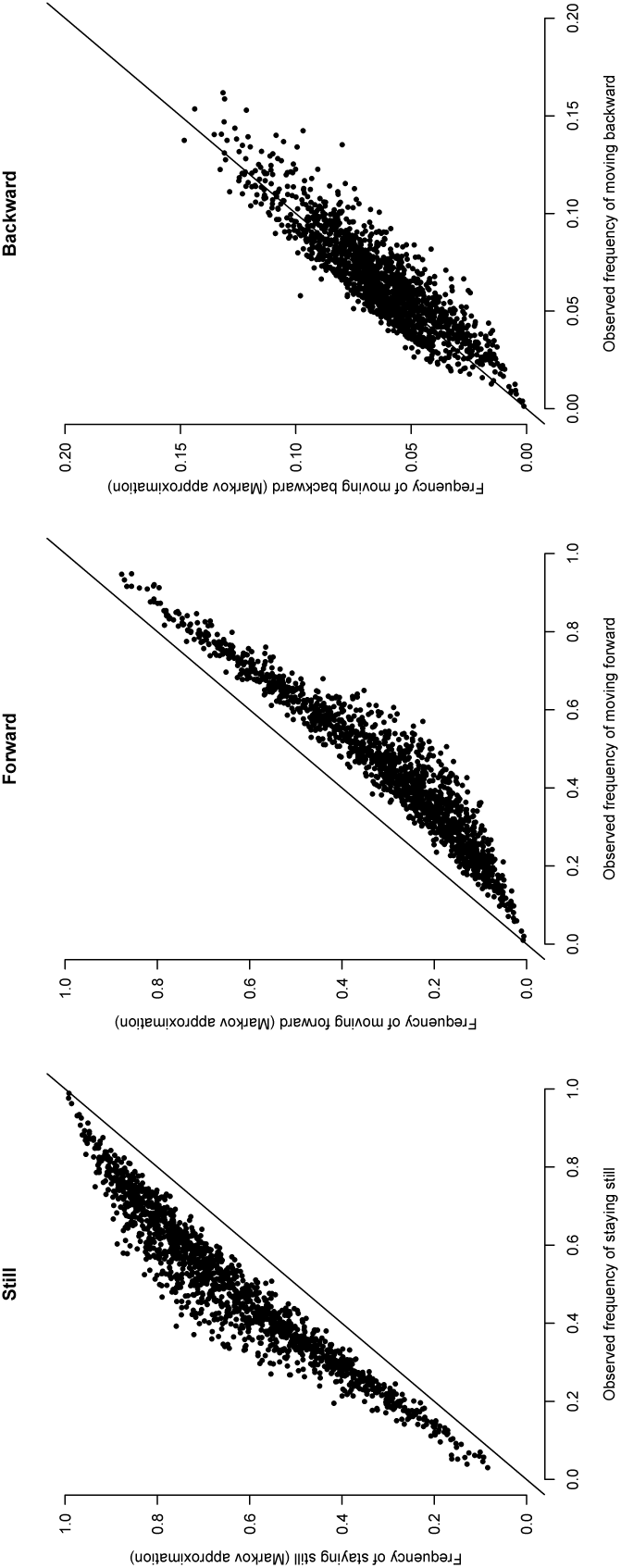
Correlation between the observed frequencies of each of the three movement states and the predicted values from the Markov model. There is a consistent bias in the long term predictions of our Markov model due to the memoryless assumption of the model being partially wrong. Some moving worms tend to remain in this state longer than expected on the long term, that is, they can be briefly interrupted but are more likely to resume movement than predicted.

**Figure S4:**
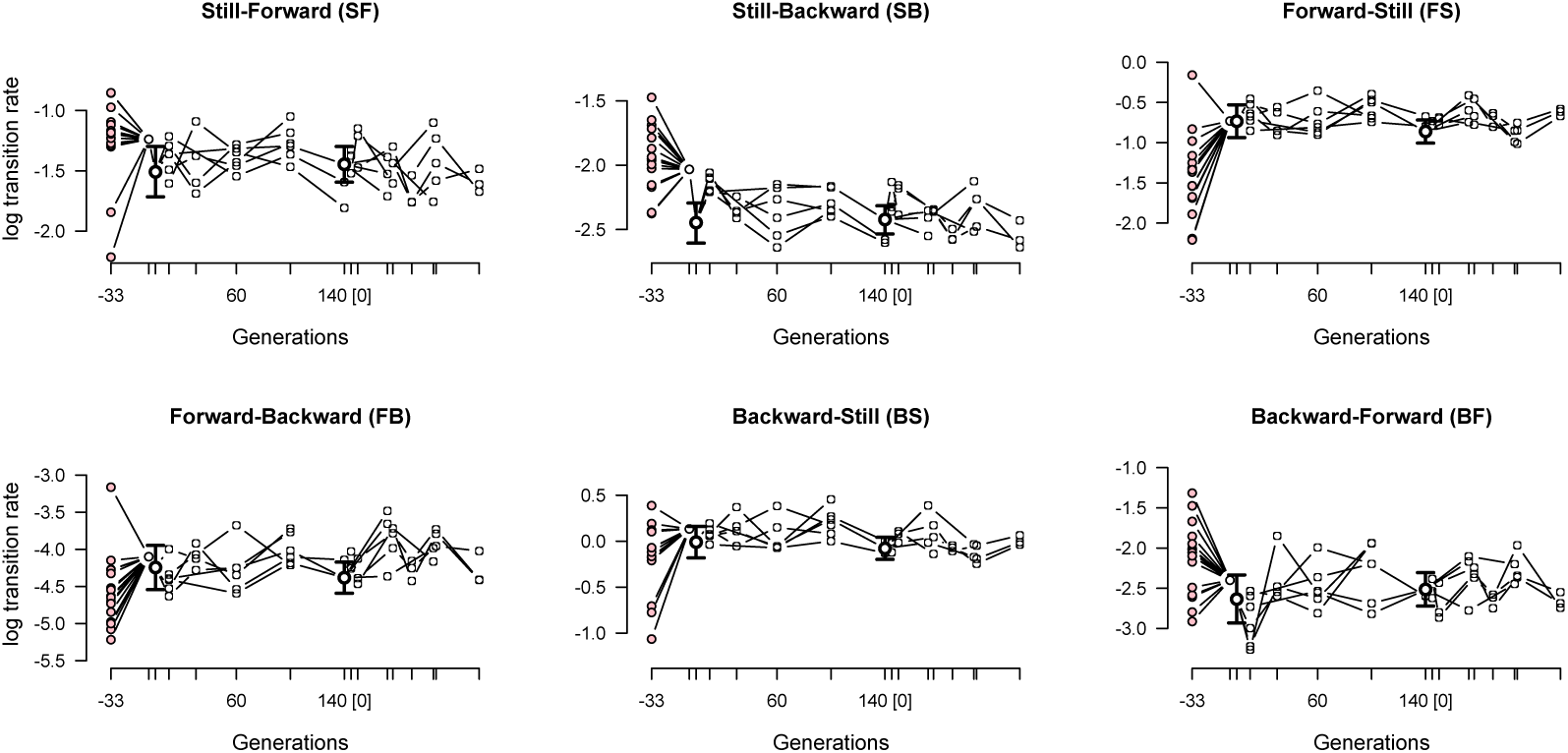
Evolution of mean hermaphrodite transition rates. Each panel shows the evolution of a transition rate in the founders (pink dots) and during experimental evolution (white dots), as shown for locomotion bias (still frequency) in Figure 1. At the beginning of the domestication and focal stages there was one ancestral population, shown by empty circles with 95% confidence intervals, while 3-6 replicate populations were measured at each sampled time point indicated by tick marks.

**Figure S5:**
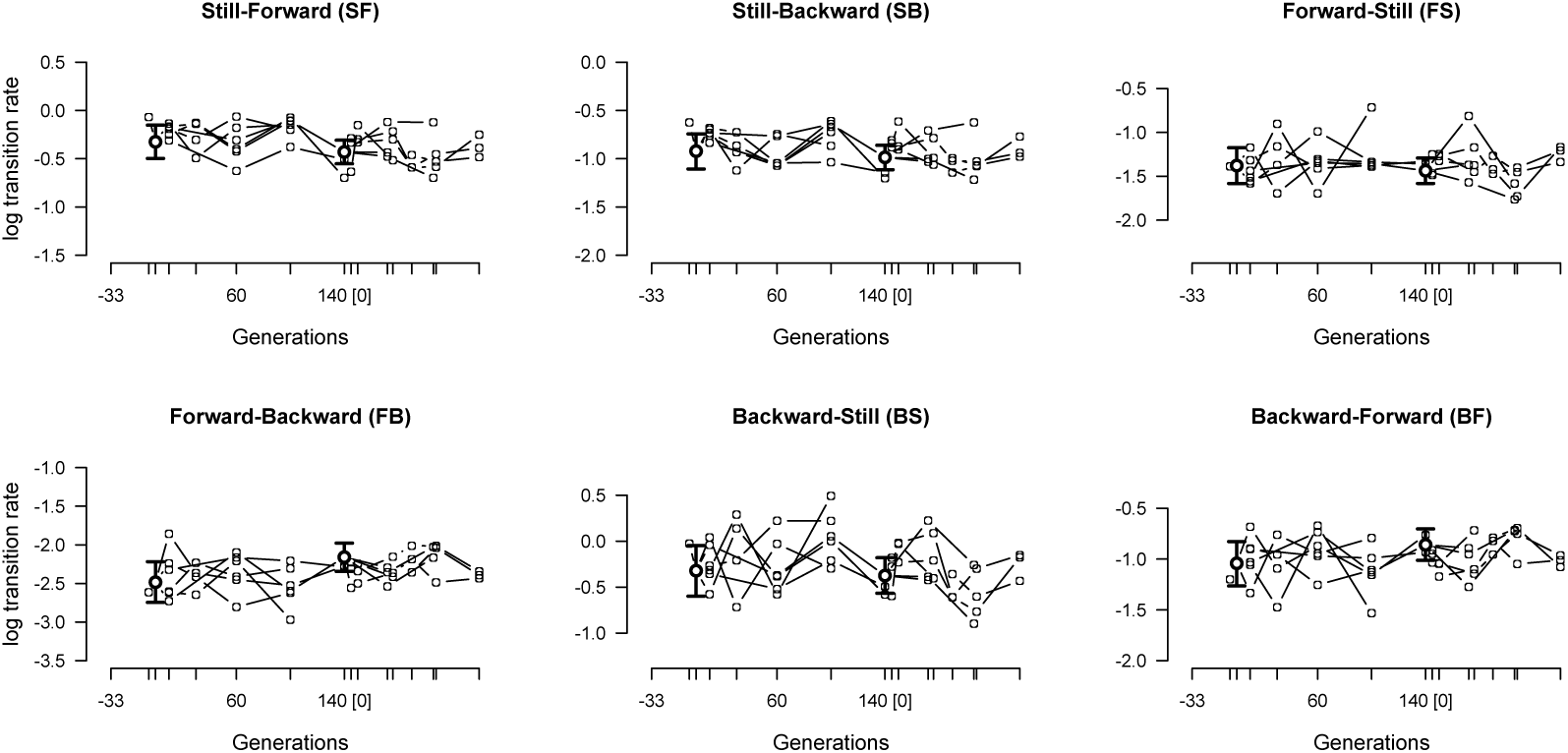
Transition rates for males (as in Figure S4).

**Figure S6:**
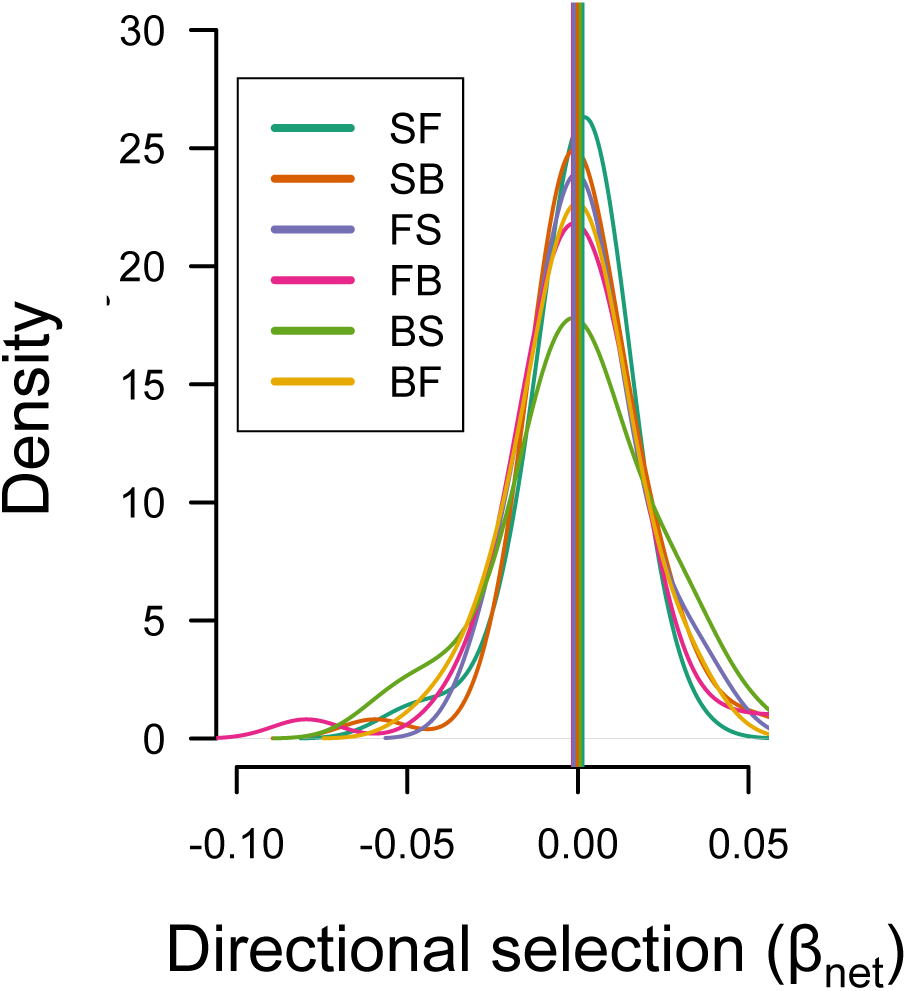
Transition rates show stasis independently of sex (Figures S4 and S5). For males, retrospective analysis of mean transition rates assuming that the **G**-matrix is stable during the period considered indicates little or no directional selection, as for hermaphrodites (Figure 2).

**Figure S7:**
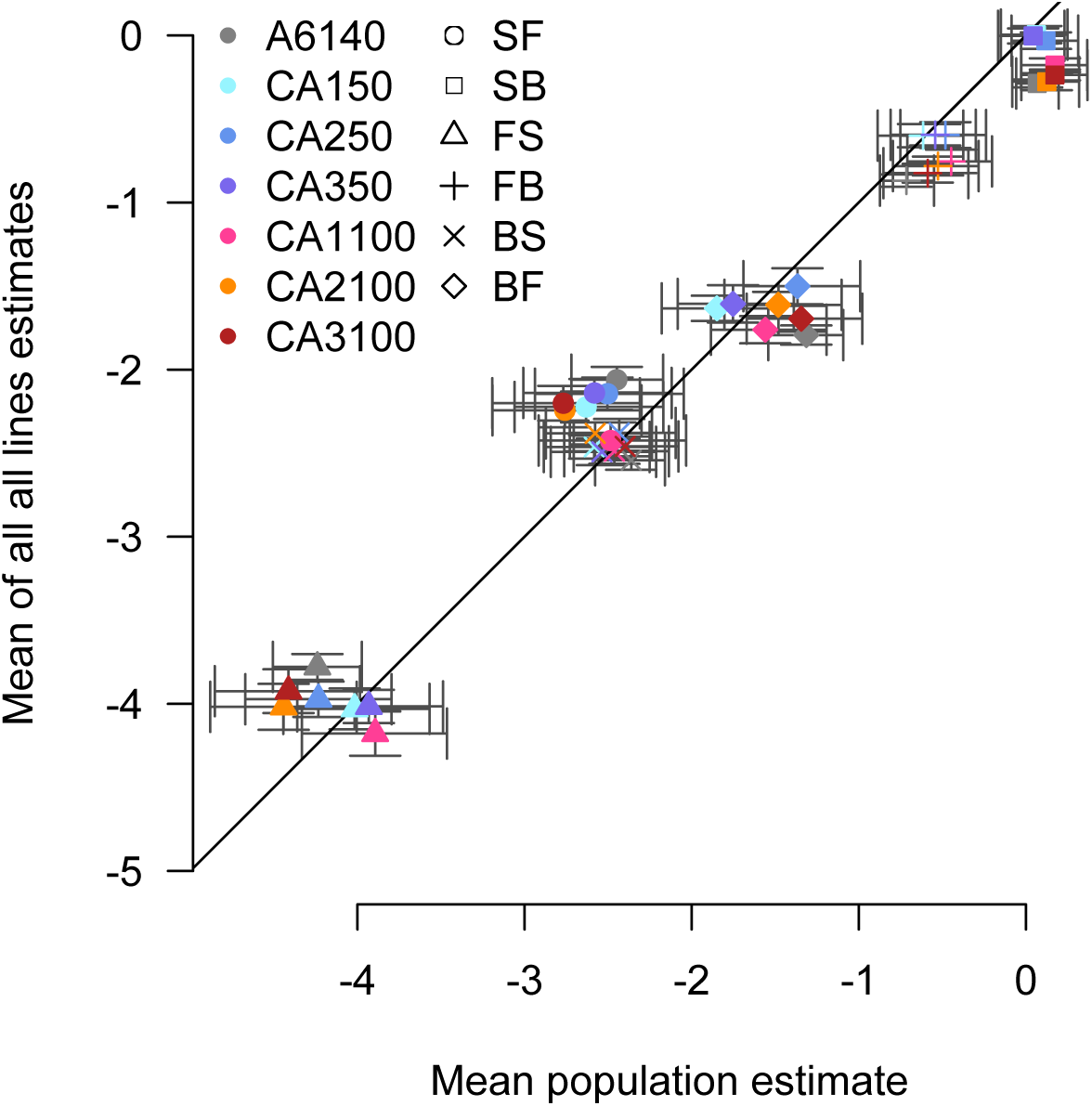
Inbreeding effects on transition rates. Mean phenotypes of all measured inbred lines show little to no significant deviations from those of their respective outbred populations. Error bars show 95 credible intervals.

**Figure S8:**
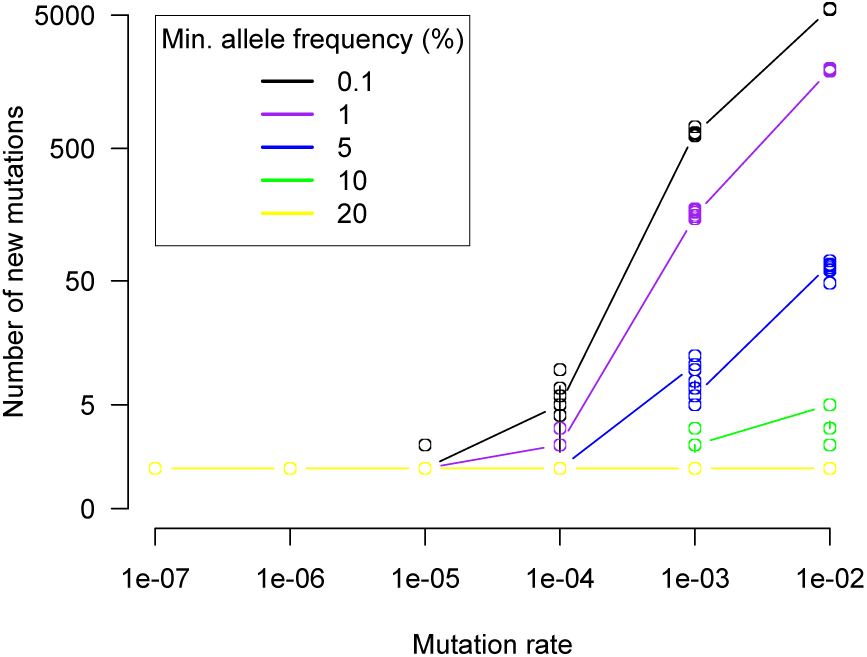
The number of new mutations reaching a given frequency threshold after 100 generations of drift. These simulations consider a fixed number (100) of freely recombining loci while controlling the probability that each loci is mutated at each generation (x-axis; equivalent to changing the mutational target size). In *C.elegans*, given a per-site SNP mutation rate on the order of 10*^−^*^9^ (Saxena et al. (2018)), a mutation rate of 10*^−^*^4^-10*^−^*^3^ would correspond to a total target size of between 1Mb and 10Mb (for reference, the average gene length is around 5Kb). Even when considering such large loci, the number of mutations reaching 5% frequency in the population is small. The contribution of neutral mutations to the observed changes in **G**-matrices is expected to be weak.

**Figure S9:**
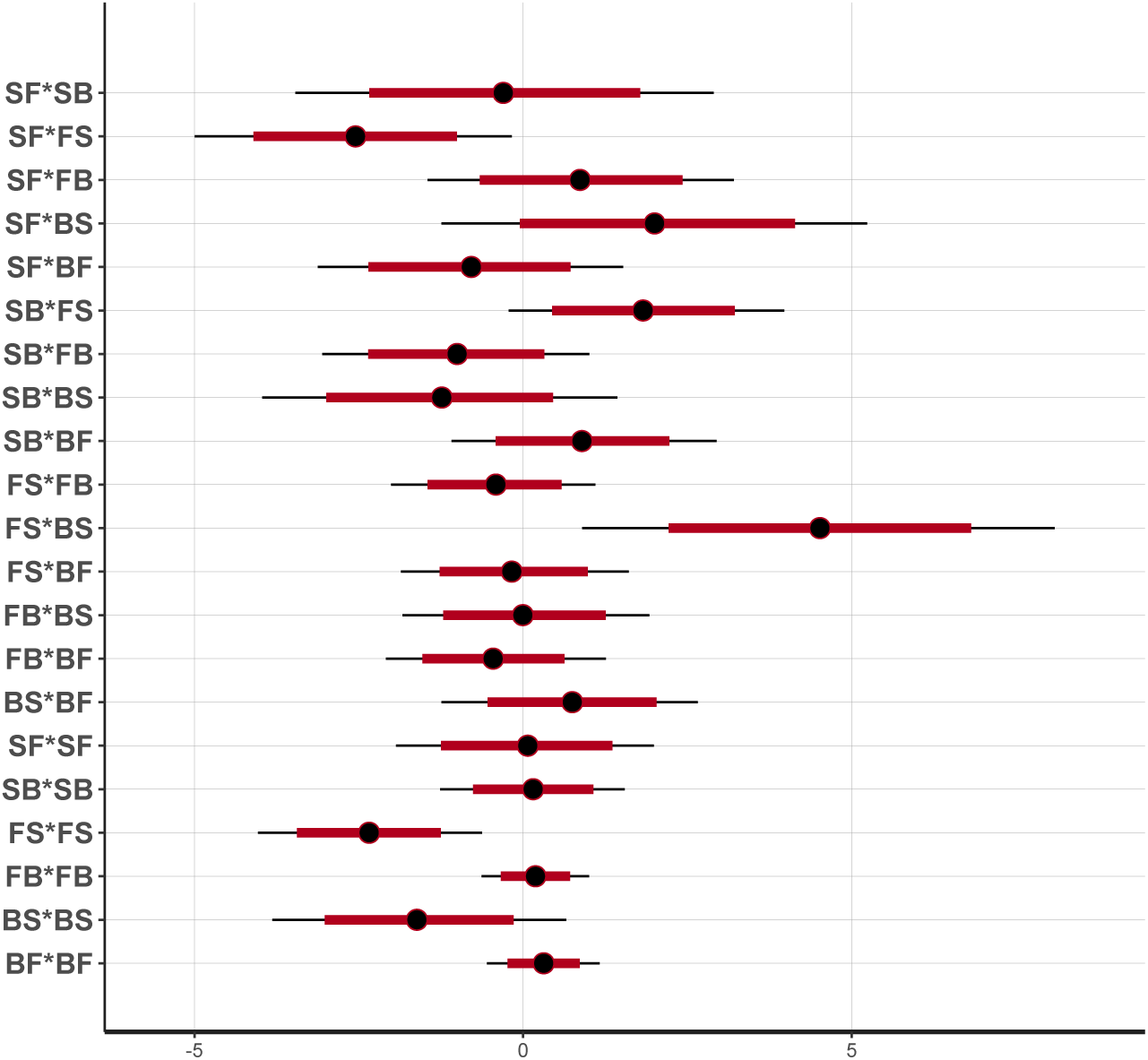
Quadratic selection coefficients. The partial regression coefficients of fertility on transition rates were estimated by Bayesian inference (Methods 5.4.2). Each row shows the median estimate (circle), and 80% and 95% credible intervals (red bar and line, respectively) of the posterior distributions. The top 15 rows show coefficients of correlated selection between two transition rates, the bottom 6 rows show coefficients of stabilizing or disruptive selection on each transition rate.

**Figure S10:**
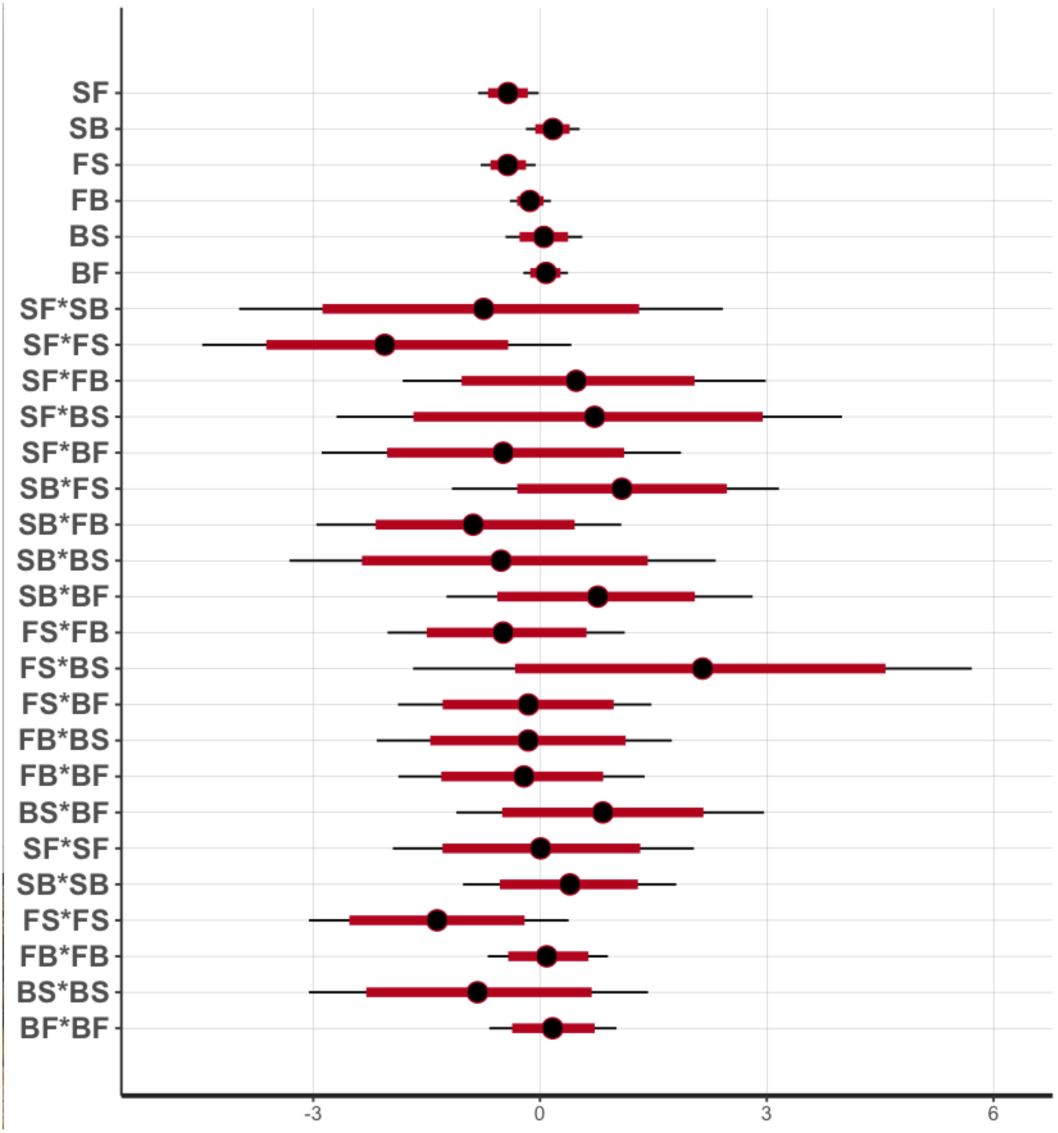
Linear and quadratic selection coefficients. As for Figure S9, with estimates from a model including linear coefficients on each transition rates (first six rows).

**Figure S11:**
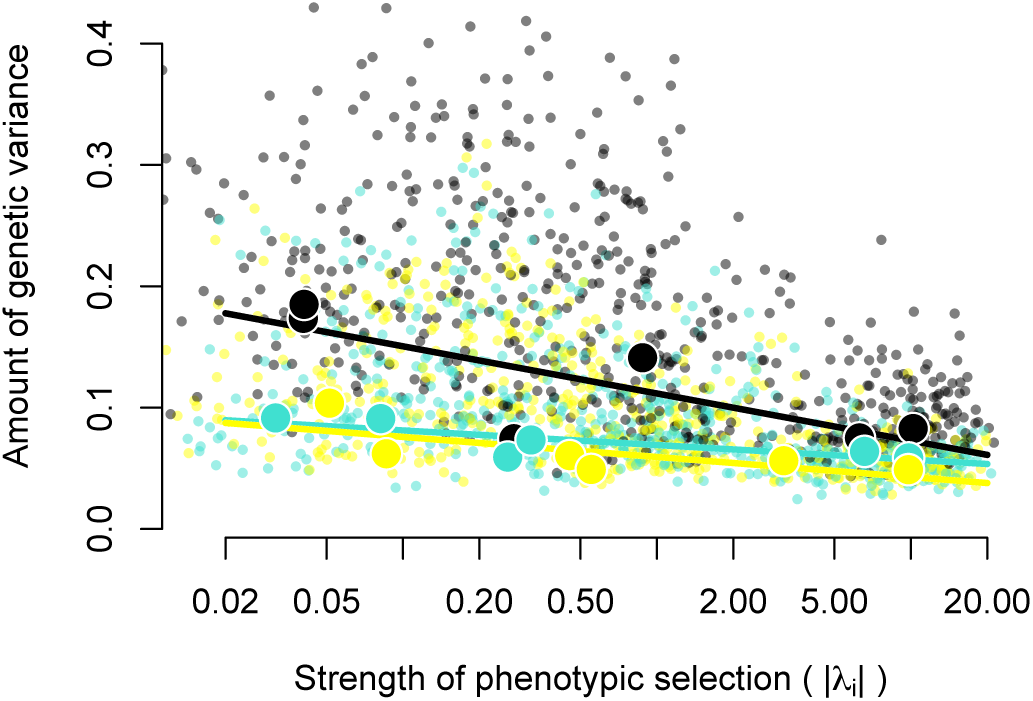
Founder and NIL projections onto the selection surface. Dots and lines were calculated as for Figure 8. The genetic variance among all 16 founders of the experimental populations are shown in black; for wild founders only in cyan; and for the CX12311 NIL (with ancestral alleles of *npr-1* and *glb-5* introgressed into the N2 background) in yellow.

**Figure S12:**
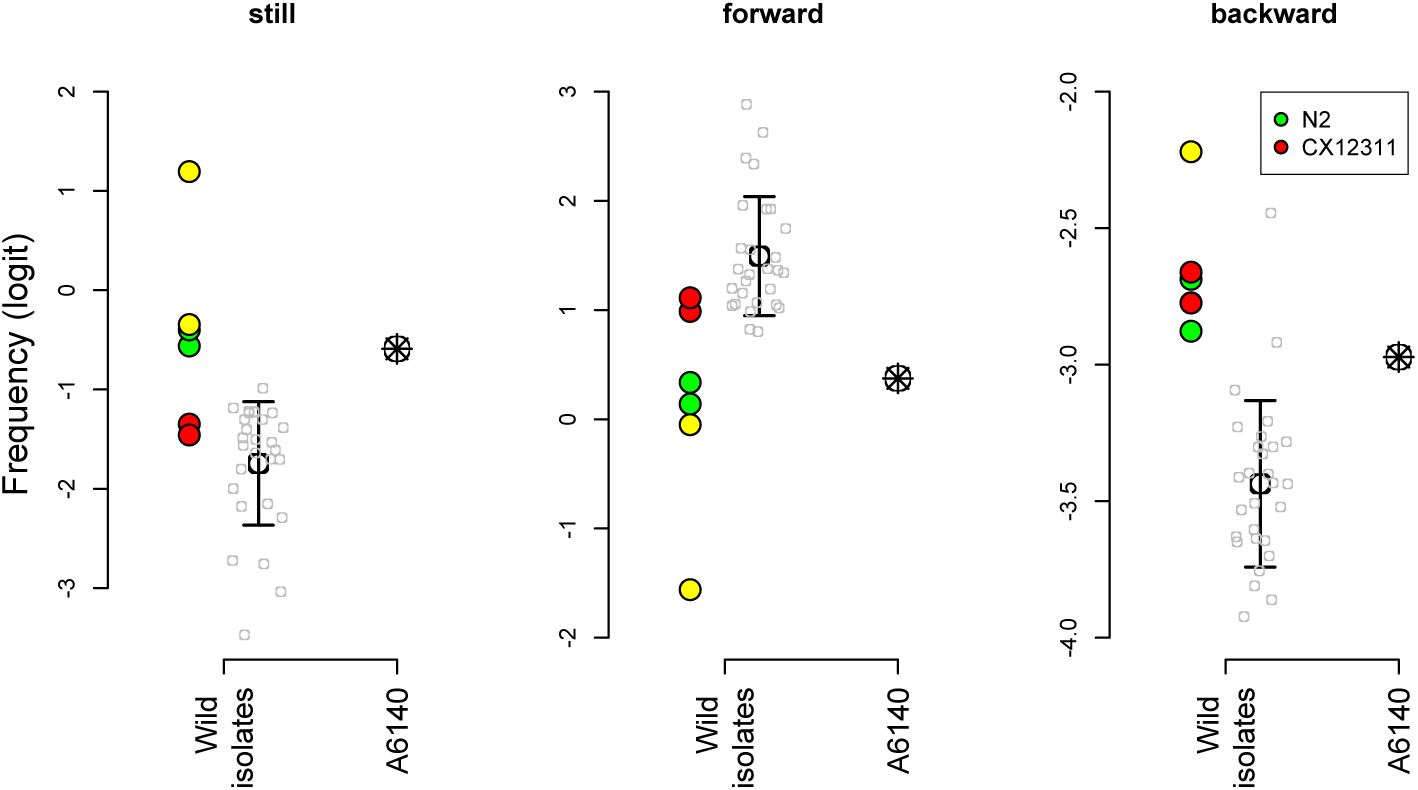
Mean frequency of each of the three locomotion states (still, forward and backward) for wild isolates (empty grey points with mean ± standard deviation); the two N2-like founders (N2 and CB4507; in green and yellow respectively); the CX12311 NIL (red), and the A6140 population (open symbol with a star). N2-like founders show a large deviation from other founders, in the direction of the rapid evolution observed during founder hybridization. Ancestral alleles of *npr-1* and *glb-5* in the CX12311 NIL explain a large proportion of this deviation for still and forward frequencies.

**Figure S13:**
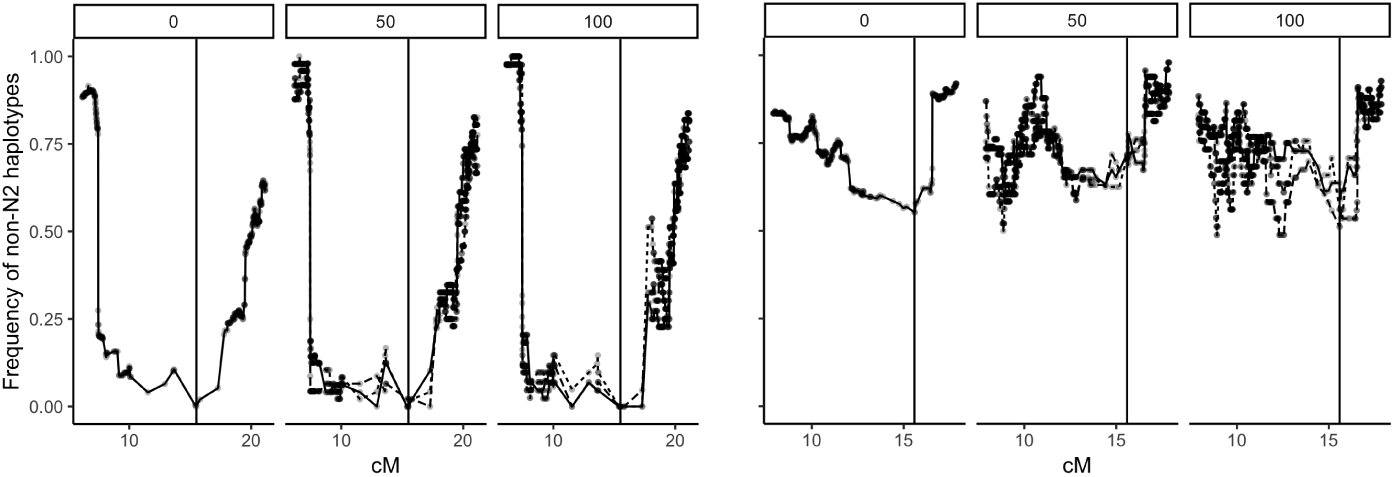
Mean frequency of wild vs. N2 haplotypes at *npr-1* (left; X chromosome) and *glb-5* (right; chromosome V) at generation 0 (A6140), 50 (CA[1-3]50) and 100 (CA[1-3]100) of the focal stage. Loci, with N2-like haplotypes at an initial frequency of 1*/*8 in the founders, are indicated with a vertical black line. The data are consistent with complete and partial sweeps for *npr-1* and *glb-5*, respectively, between hybridization and sampling in the A6140 after domestication. However, *glb-5* has been maintained at intermediate frequencies over the following 100 generations in the six replicate populations. Haplotypes were reconstructed from founder and RIL genotypes (Noble et al., 2019) conditioned on N2/CB4856 recombination data (Rockman and Kruglyak, 2009), and scaled to an *F*_2_ chromosome length of 50cM. Points are plotted at SNP markers subsampled to 0.01cM resolution.

**Figure S14:**
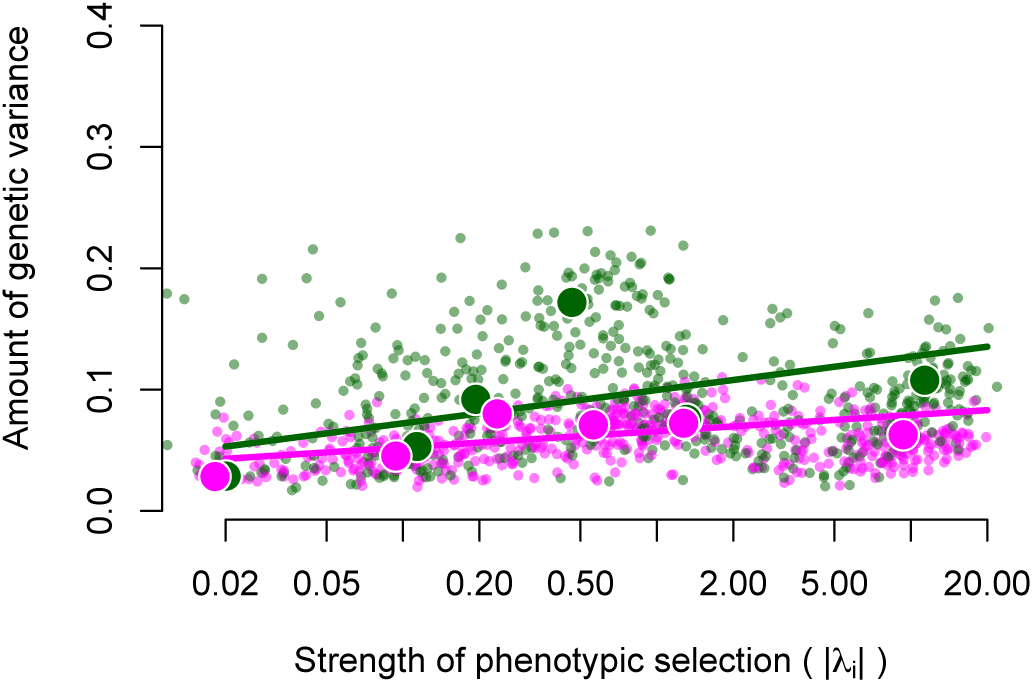
MA line projections onto the selection surface (N2 background in dark green, PB306 background in purple). Dots and regression lines were calculated as for Figure 8.

**Figure S15:**
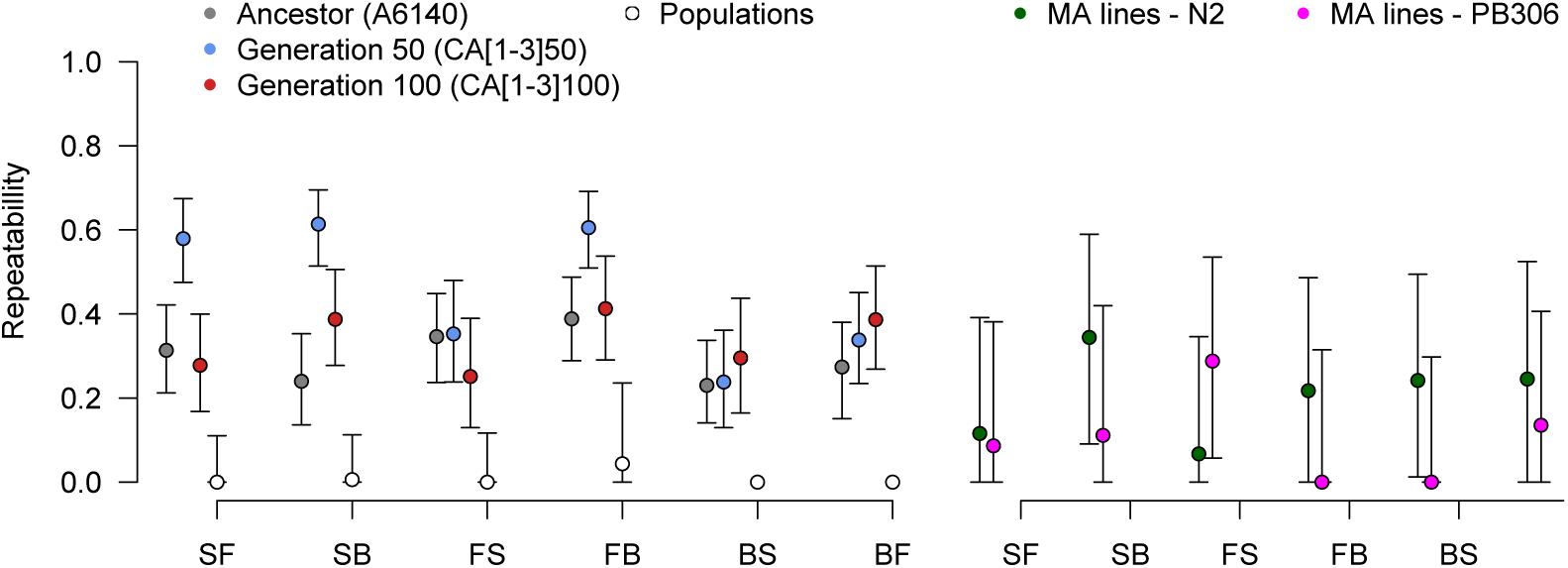
Repeatability of transition rates in hermaphrodites. Left panel: inbred lines from the focal stage and all outbred populations. Right panel: MA lines in N2 and PB306 backgrounds. Error bars are 95% bootstrap confidence intervals.

**Figure S16:**
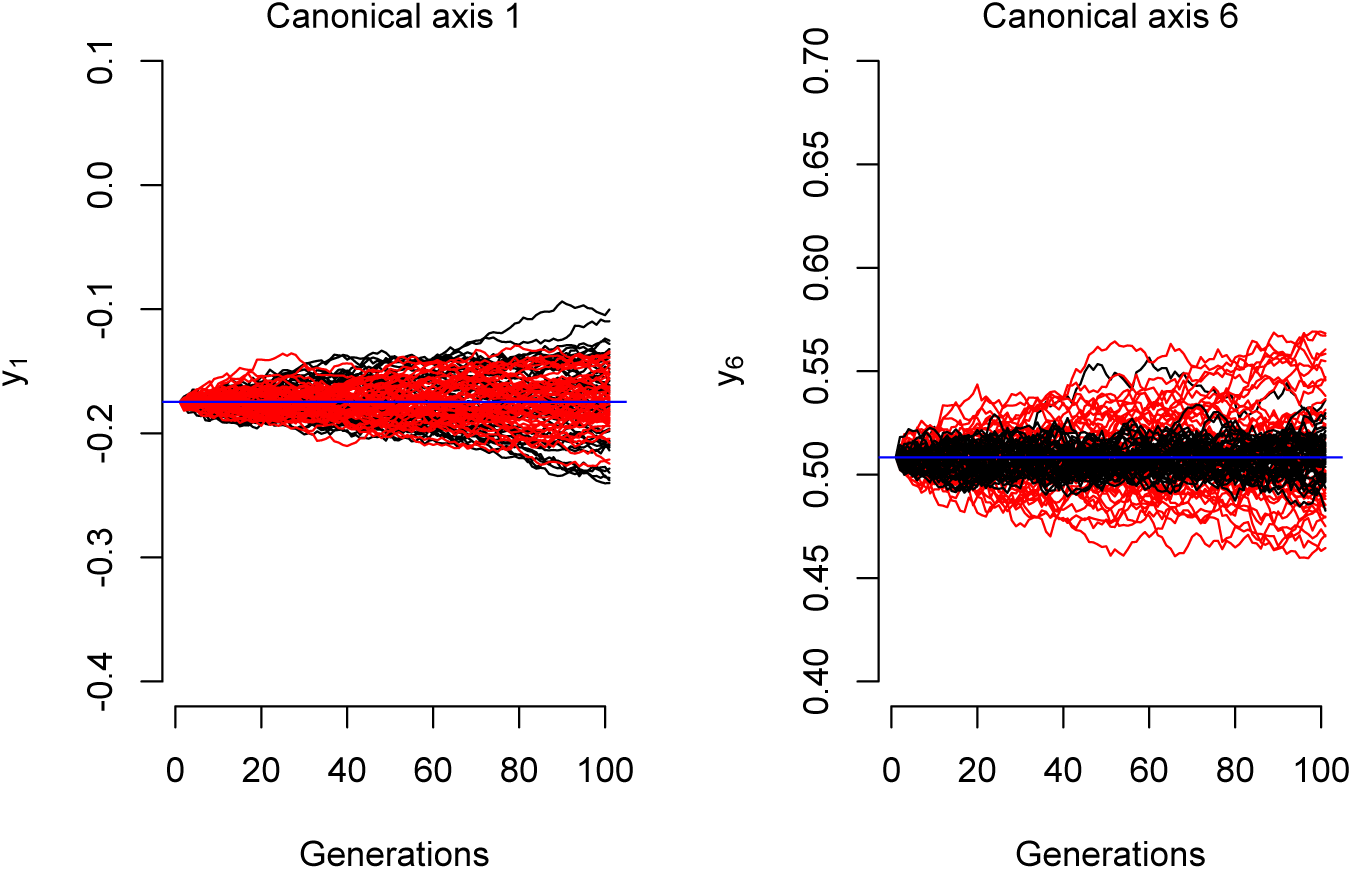
Phenotypic divergence among 50 replicate simulations of an OU process (black) or drift (Brownian motion, red) along the canonical axes with the strongest disrupting and stabilizing selection in our environment (*y*_1_ and *y*_6_, see Figure 7). The **G**-matrix was assumed to be that of the ancestral domesticated A6140 population, with *N_e_* = 10^3^ (Chelo and Teotónio, 2013). Disruptive selection leads to more divergence than expected by drift alone, while stabilizing selection maintains the mean trait value of the population near the phenotypic optimum.

**Figure S17:**
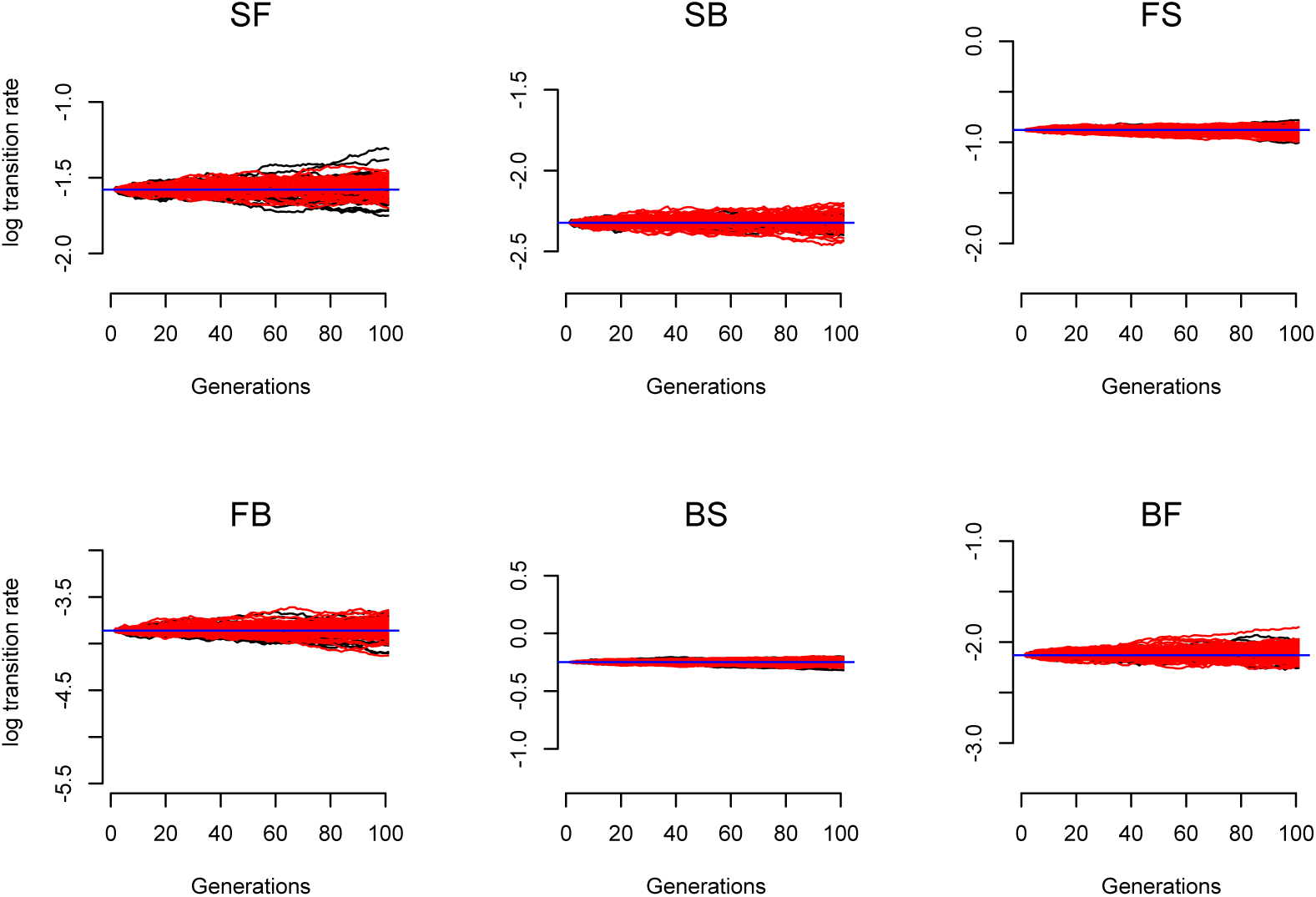
Divergence of transition rates, simulated as in Figure S16. There is no clear signal of stabilizing or disruptive selection on any of the traits. Divergence from the ancestral state is apparent and thus the OU does not predict phenotypic stasis, as observed during experimental evolution (see also main Figure 9).

## 8 Supplementary Table

**Table S1:**
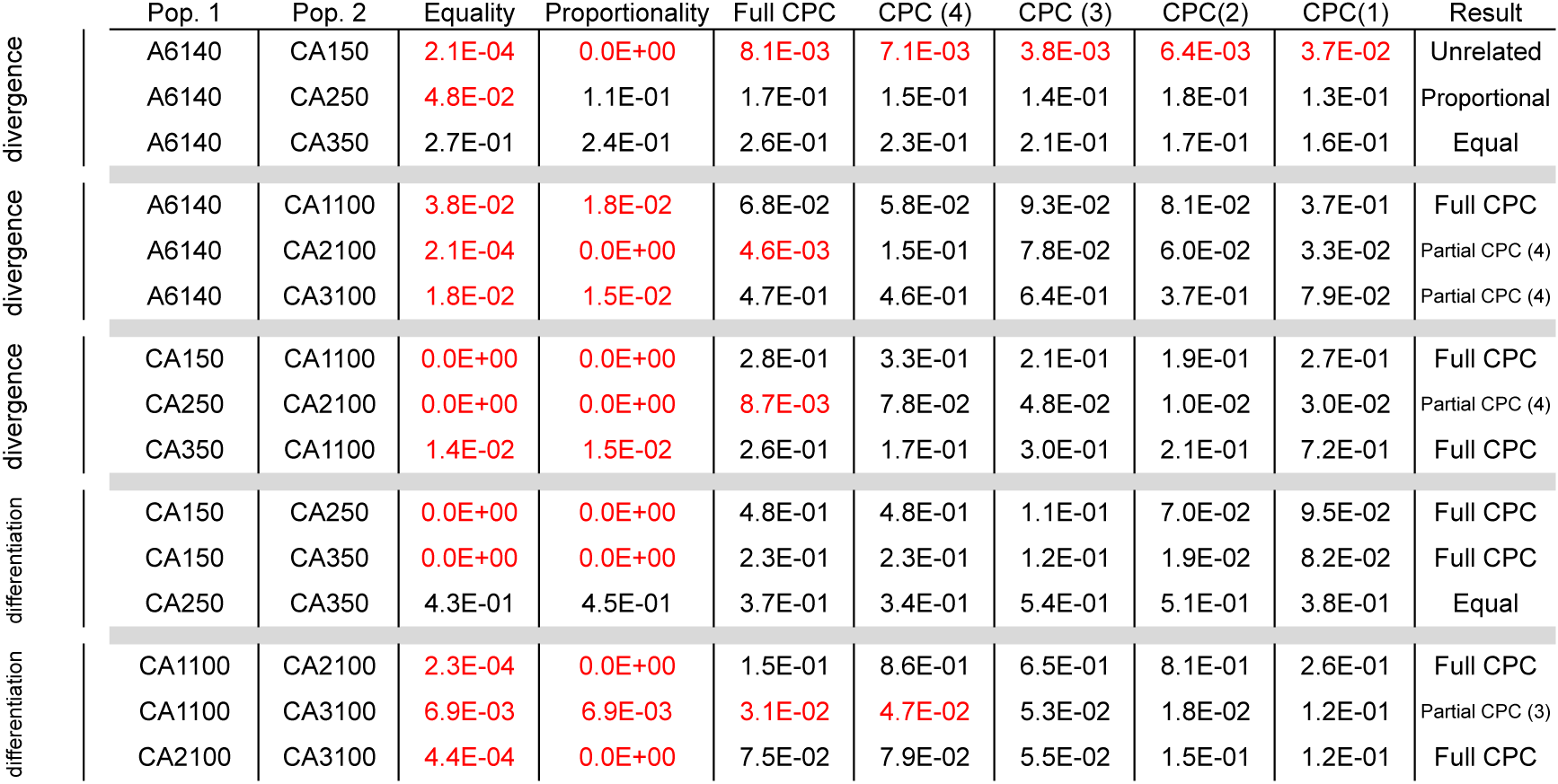
Divergence and differenciation of the G-matrices during experimental evolution. The table present the results of the Flury comparison that tests a succession of hierarchichal hypothesis concerning the degree of similitude between two matrices. At each degree of similitude in the hierarchy, a hypothesis is tested against the hypothesis at the bottom of the hierarchy (i.e., unrelated). Starting from the test of equality, if the test is rejected, one should consider a lower degree of similitude, and so on.The first two columns present the two matrices tested. The following 7 columns present the p-values of the hierarchical hypothesis from top to bottom: equality, proportionality, all principal components being shared (“Full CPC”) or only a subset of them (CPC (#). Significant tests are highlighted in red. The last column present the conclusion of the method, matching the first non-significant test in the hierarchy.

## 9 Archiving

Data and code for analysis are listed as separate files, archived in our github page (https://lukemn.github.io/cemee/), and will be archived in XXXX upon publication:

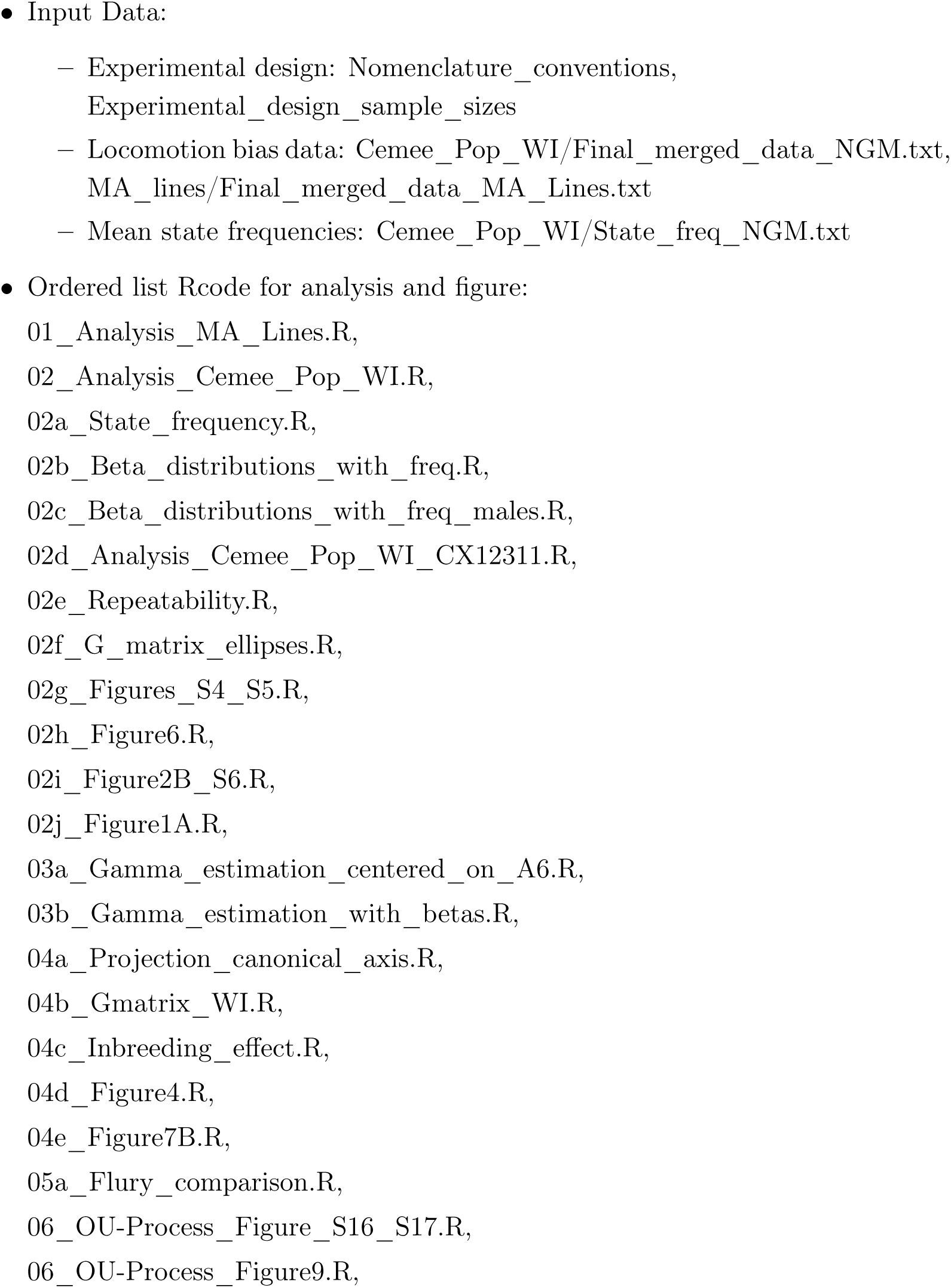
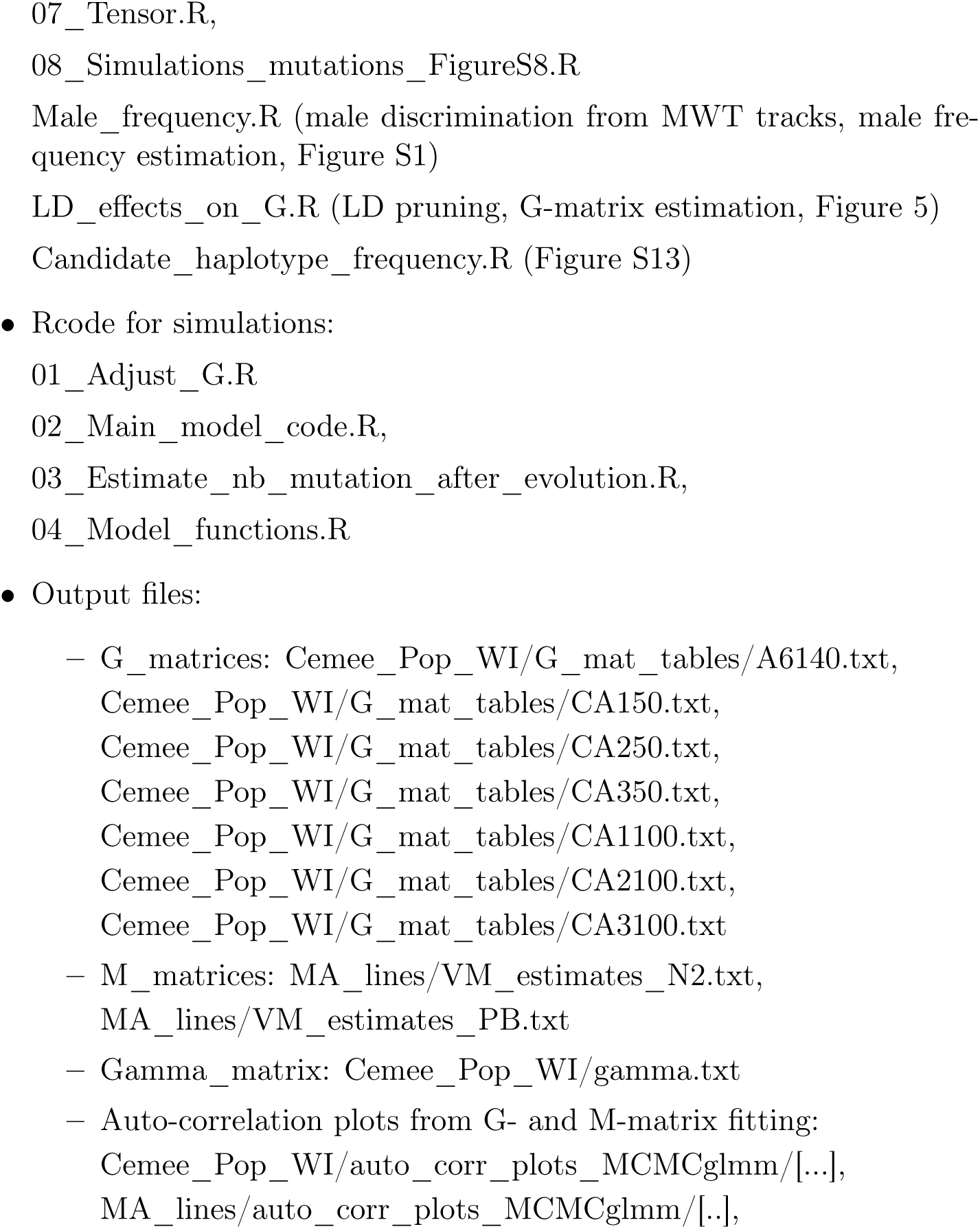

## 10 Acknowledgements

We thank A. Crist, J. Garcia, H. Gendrot, C. Goy, V. Pereira, F. Melo, and A. Silva for help with worm handling and data acquisition; P. McGrath for worm strains; R. Costa, R. Kerr, S. Proulx, P. Phillips, N. Scwierczek for help with data analysis, hardware and software implementation; L.-M. Chevin, C. Dillmann, M.-A. Félix, P. Phillips, A. Le Rouzic, and A. Veber for discussion.

## 11 Funding

This work was supported by the European Research Council (ERC-St-243285) and the Agence Nationale pour la Recherche (ANR-14-ACHN-0032-01, ANR-17-CE02-0017-01) to HT, and the National Institutes of Health (R01GM107227) to CB. BA was from 2010-2015 a Fundação para a Ciência e a Tecnologia PhD scholarship recipient (SFRH/BD/51177/2010). LN is a Marie Curie fellow (H2020-MSCA-IF-2017-798083).

## 12 Competing interests

We have no financial or non-financial interests to declare.

## 13 Author contributions

Conceptualization FM, LN, CB, HT; hardware and software implementation BA, FM, TG; data acquisition and analysis FM, LN, BA, TG; funding acquisition LN, CB, HT; project administration HT; resources CB, HT; writing, original draft FM, HT; writing, review and editing LN, CB; correspondence FM, LN, HT.

## Notes

https://lukemn.github.io/cemee/

